# BCG vaccination elicits protection against Mtb infection mediated by two phases of T cell immunity

**DOI:** 10.1101/2025.10.28.685186

**Authors:** Abiola F. Ogunsola, Rocky Lai, Kelly Cavallo, Gillian L. Beamer, Samuel M. Behar

## Abstract

Vaccine development for tuberculosis is a global priority. Our studies using Collaborative Cross (CC) mice show that genetic diversity influences the efficacy of BCG, the most widely used TB vaccine. BCG vaccination of CC042 mice reduces their lung bacillary burden and increases their survival following low-dose aerosol *Mycobacterium tuberculosis* infection (MTBI), despite impaired T cell trafficking from a defective *Itgal* gene. Early protection requires the presence of T cells at the time of BCG vaccination but is not mediated by B cell or T cell recall responses following MTBI. In contrast, T cell depletion following BCG vaccination reduces survival after MTBI. Thus, CC042 mice reveal two phases of immunity induced by BCG that require T cells: an early phase mediated by innate responses and a later phase mediated by effector CD4 and/or CD8 T cells. Although measurement of vaccine-induced protection 30 days after MTBI is a standard measure of vaccine efficacy in the murine TB model, we find this time point is independent of memory T cells. Our results suggest that vaccine-elicited innate responses have a larger role in protection than previously considered. The concordance between lung CFU, pathology, and survival make CC042 mice a useful model for vaccine evaluation.

## Introduction

Tuberculosis (TB) is a chronic bacterial disease that most commonly affects the lungs. It is caused by infection with the intracellular pathogen *Mycobacterium tuberculosis* (Mtb), and is the leading cause of death worldwide by a single infectious agent (1). The only approved vaccine for TB prevention is BCG, a live-attenuated strain of *Mycobacterium bovis* (2). BCG prevents disseminated disease, such as miliary and meningeal TB in infants. However, BCG’s efficacy against pulmonary TB in adolescents and adults, which is the contagious form of the disease and drives the global TB epidemic, ranges from 0% to 80% across different populations (3). Despite BCG’s widespread use in endemic countries for nearly a century, this single vaccine has not reduced the number of new cases (8-10 million) reported each year (1), highlighting the need for an improved understanding of how BCG does and does not induce protection.

The variation in BCG’s efficacy is attributed to several factors including geography and climate, BCG strain, and age at initial vaccination (3–6). The impact of host genetic variation on BCG’s ability to confer protection against TB has gained recent interest but open questions remain (7). The murine TB model has relied on one or two genetically related inbred mouse strains for most experimental studies. While studies in C57BL/6 (B6) and other inbred strains have been invaluable in defining the immunological response to infection, there has yet to be a successful vaccine developed using the mouse model that has been effective in people. Although it is uncertain whether this shortcoming is a problem with the model or the vaccines, the reliance on inbred mice constrains our understanding of how host genetic variation modifies susceptibility to infection and shapes immune responses stimulated by vaccines.

To facilitate identifying correlates of vaccine-induced protection that could be applicable to diverse human populations (8), we turned to the Collaborative Cross mouse resource. The Collaborative Cross (CC) is a series of recombinant inbred mouse strains derived from eight genetically distinct parental strains, which was developed for the purpose of incorporating genetic analyses into experimental studies using mice (9–11). There is a wide variation in susceptibility to Mtb infection among the different CC mouse strains (12, 13). BCG-mediated protection among CC strains also varies with genetic background, and vaccine-induced protection is a trait that appears to be distinct from Mtb susceptibility (10). We find that BCG elicits a variety of immune responses among CC mice, highlighting the need to identify vaccine-induced correlates of protection through the lens of genetic diversity (14).

CC042 mice are highly susceptible to Mtb, and rapid TB disease progression has been attributed several genetic loci. One of these is a splice variant in the *Itgal* gene that results in a truncated, defective protein, and reduced capacity to recruit lymphocytes to the lungs following infection (15). *Itgal* encodes CD11a, the alpha chain (αL) of the heterodimeric αLβ2 integrin known as <u>L</u>eukocyte <u>F</u>unctional associated <u>A</u>ntigen-1 (LFA-1) (16). LFA-1 is an important adhesion molecule expressed on the surface of lymphocytes and it contributes to immune cell trafficking and immune synapse formation (16–18). Similar to CC042 inbred mice with mutated *Itgal,* B6 mice completely lacking the αL integrin (i.e., CD11a*^−/–^* mice) are also more susceptible to Mtb, further indicating that LFA-1-mediated T cell trafficking to the lungs is critical for Mtb control (19).

Here we report that despite the genetic mutation in *Itgal* leading to dysfunctional LFA-1 and poor T cell trafficking to the lungs (10), BCG vaccination protects CC042 mice against low-dose aerosol Mtb infection. Given the defect in T cell recruitment to the lung, we hypothesized that BCG elicits protection in CC042 mice by a mechanism independent of memory T cells. Such a mechanism could be unique to CC042 mice, or the genetic mutation in *Itgal* in CC042 mice might unmask important immune mechanisms of vaccine-mediated protection. Thus, we surmised that understanding the mechanism of BCG-mediated protection in CC042 mice would reveal mechanisms of vaccine-induced immunity independent of LFA-1. We find that lymphocyte trafficking to the lung is not necessary for BCG-mediated protection of CC042 mice, early (i.e., 4 weeks) after MTBI. In fact, deletion of memory T cell and B cell responses are dispensable for short-term protection against Mtb. However, durable protection (e.g., survival), requires memory T cells. Thus, CC042 mice reveal that BCG-induced protection against low dose aerosol Mtb infection at the standard time point of four weeks requires T cells to be present at the time of vaccination but is not mediated by conventional CD4 or CD8 T cells, or B cells. In contrast, memory T cells are crucial for long-term survival conferred by BCG vaccination. Our results provide insight into the mechanisms of BCG-elicited immune protection and establishes CC042 mice as a new model for preclinical vaccine testing.

## Results

### BCG vaccination protects CC042 mice against Mtb challenge

To validate the protection results in our screen (10), cohorts of B6 and CC042 mice vaccinated subcutaneously (s.c.) with BCG were compared to unvaccinated mice. After 12 weeks of rest, all mice were challenged with low-dose aerosolized Mtb (strain Rv.YFP) for four weeks (Fig.1A). Following a 4-week infection, both B6 and CC042 mice are protected by BCG vaccination. The bacterial burden in the lungs, mediastinal lymph node (mLN) and spleen in both strains is significantly reduced following BCG vaccination, compared to the non-vaccinated cohort (Fig.1B-D). These results were consistent across multiple experiments (Fig.1E-G). Interestingly, the magnitude of protection conferred by BCG vaccination (the net reduction of Mtb CFU in vaccinated vs. unvaccinated mice) is greater in CC042 mice, than in B6 mice, which typically show a 10-fold (i.e., 1 log_10_) reduction in the Mtb in the lung (Fig.1H). Lastly, consistent with previous findings, the lungs of CC042 mice had significantly greater Mtb numbers compared to B6 mice, highlighting the increased susceptibility of CC042 mice (Fig.1B). The bacterial burdens in the mLN and spleen of CC042 mice were also greater than that of B6 mice but did not reach statistical significance (Fig.1C-D).

**Figure 1.**
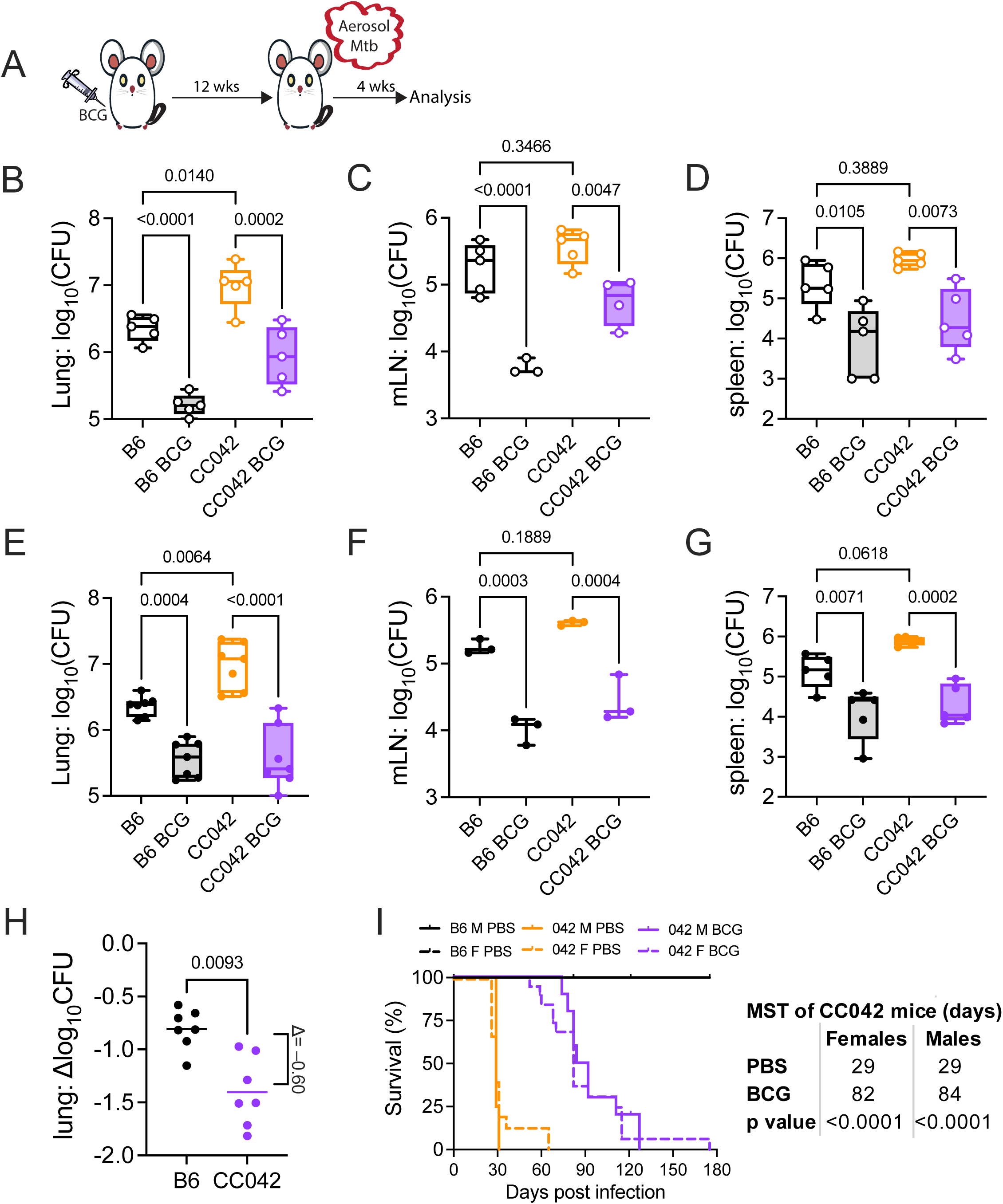
BCG protects CC042 mice against TB. A. Experimental scheme. B6 and CC042 mice were BCG vaccinated or not and rested for 12 weeks. Then, all mice were infected with Rv.YFP and analyzed after four weeks. B-D. Representative experiment showing CFU in the lung (B), mLN (C), and spleen (D), n=4-5 mice per group. Each point is an individual mouse. One-way ANOVA with Šídák’s multiple comparisons test. Bars, mean. Error bars, SD. E-G. Results combined from seven (E, lung), three (F, mLN), and five (G, spleen) experiments. Each point is the mean of an independent experiment. One-way ANOVA with Šídák’s multiple comparisons test. Bars, mean. Error bars, SEM. H. CFU reduction in the lung from seven experiments. Δlog_10_CFU= log_10_CFU (BCG) - log_10_CFU (unvaccinated). Paired t-test. Each point represents an independent experiment. Line, mean. I. Cumulative survival of non-vaccinated B6, non-vaccinated CC042 and BCG-vaccinated mice after aerosolized Mtb Erdman infection. Data compiled from three independent experiments. 15 B6 female mice, 10 B6 male mice, 15 CC042 female mice, 12 CC042 male mice, 19 CC042 female BCG-vaccinated mice, 11 CC042 male BCG-vaccinated mice. The difference between non-vaccinated B6 and non-vaccinated CC042 and the difference between non-vaccinated CC042 and BCG vaccinated CC042 is statistically significant as determined by the Mantel-cox log-rank test (p < 0.0001).

We then asked if BCG vaccination confers a survival benefit to CC042 mice following Mtb challenge. A BCG-vaccinated cohort of CC042 mice was compared to unvaccinated CC042 and unvaccinated B6 mice. The mice were rested for 12 weeks after vaccination, and then all mice were challenged with low dose aerosolized Mtb Erdman. Following Mtb infection, unvaccinated B6 mice survived for more than 180 days while the unvaccinated CC042 mice succumbed to infection with a median survival of 29 days (Fig.1I). BCG vaccinated CC042 mice had a median survival of 82-88 days following infection. The protective effect of vaccination is independent of sex as BCG confers a significant survival benefit to both male and female CC042 mice. These data show that BCG confers durable protection to CC042 mice despite their lack of CD11a.

### BCG vaccination promotes lymphocyte infiltration into the lungs during Mtb infection

To understand how CC042 mice, which have a defect in CD11a, are protected from Mtb by BCG vaccination, the caudal lung lobes were collected at the four weeks post infection (wpi) endpoint (Fig.1A), fixed in formalin, and using a acid-fast bacteria stain (AFB) in combination with hematoxylin and eosin (H&E), stained to assess the composition of cellular infiltrates (Fig.2). Four weeks after infection, unvaccinated B6 mice, have mild to moderate disease with more circumscribed lesions containing well-defined lymphocytic infiltrates, few AFB that are mostly single bacilli, although clusters are also identified (Fig.2, first row). BCG-vaccinated B6 mice have mild disease with prominent lymphocyte cuffing and dense lymphocytic infiltrates, few foamy macrophages with small foci of neutrophils and rare AFBs (Fig.2, second row). In contrast, CC042 mice have mild to moderate disease due to diffusely organized lymphohistiocytic lesions with AFB frequently observed in clusters (Fig.2, third row). The lesions in BCG-vaccinated CC042 mice are similar to the B6 lesions: they are smaller and well-defined, containing more lymphocytic infiltrates than unvaccinated mice and fewer AFB (Fig.2, fourth row). Overall, in the unvaccinated state, susceptible CC042 mice have disorganized, loosely structured, lymphocyte-poor lung lesions and poor control of Mtb replication, compared to inherently resistant B6 mice. BCG vaccination of CC042 mice leads to a more compact lesion with more lymphocytes and fewer bacilli.

**Figure 2.**
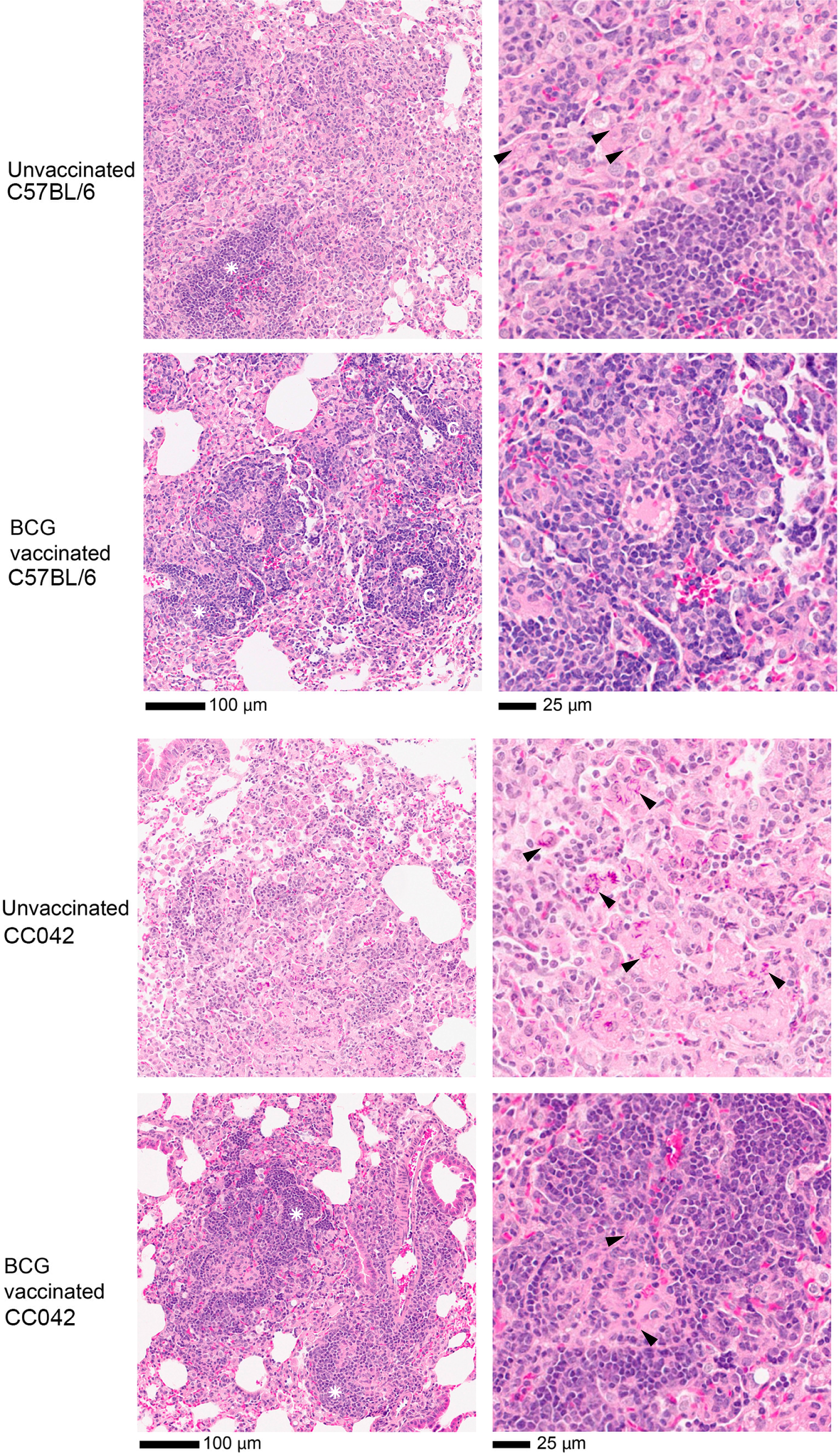
BCG reduces the severity of disease in the lungs of CC042 mice. The caudal lung lobe from unvaccinated or BCG vaccinated mice were obtained four weeks after Mtb challenge, and sections were stained with a combined AFB and H&E stain. The 1^st^ and 2^nd^ row are images from unvaccinated or BCG-vaccinated B6 mice, respectively. The 3^rd^ and 4^th^ row are images from unvaccinated or BCG-vaccinated CC042 mice, respectively. Asterisks indicate AFB, although not all AFB are designated. Scale bars are shown for reference. Each image is representative of an individual subject (n=4-5/group), and the images are from one of three independent experiments with similar results. Black arrowheads, AFB; White asterisks, lymphocyte infiltrates; ‘C,’ lymphocyte cuffs.

### BCG-mediated protection of CC042 mice does not require lymphocyte trafficking to the lung

BCG vaccination and Mtb infection stimulates immune responses that result in tissue resident T cells in both humans and mouse models (20, 21). We hypothesized that BCG vaccination of CC042 mice elicits T cells that traffic to the lung independently of CD11a and become lung resident memory T cells (i.e., TRM) (18, 22, 23). To completely disrupt trafficking of lymphocytes to the lung, we used the drug FTY720, which degrades the sphingosine-1-phosphate receptor (S1P-R) and prevents lymphocyte egress from lymph nodes (LN) (24). The conditions for administration of FTY720 were optimized in B6 mice. We found that while both low (1 mg/kg) and high (4 mg/kg) doses of FTY720 successfully inhibit lymphocyte trafficking to the lung, only high dose FTY720 led to a statistically significant increase in the bacterial burden in the lungs of B6 mice (Fig.S1). A dose of 4 mg/kg was used in all subsequent experiments.

Groups of CC042 and B6 mice were vaccinated with BCG, while control groups were left unvaccinated. After 12 weeks, all mice were treated with FTY720 commencing the day before low-dose aerosolized Mtb Rv.YFP infection (Fig.3A). FTY720 treatment reduced circulating T and B cells during the 4-week infection period in both unvaccinated and vaccinated CC042 mice by 83-93% (Fig.3B, and Fig.S2 for gating scheme). Next, the Mtb burden in the lungs and spleens of all mice was determined. FTY720 treatment of unvaccinated B6 mice led to an increase in Mtb CFU in the lung and spleen (Fig.3C). In contrast, FTY720 treatment of unvaccinated CC042 mice had no effect on the bacterial burdens in the lung and spleen (Fig.3D). In BCG vaccinated B6 and CC042 mice, protection, as measured by lung and spleen CFU, was maintained despite FTY720 treatment. Thus, BCG-mediated protection of C0042 mice is fully maintained even when lymphocyte circulation and trafficking to tissues is almost completely inhibited.

**Figure 3.**
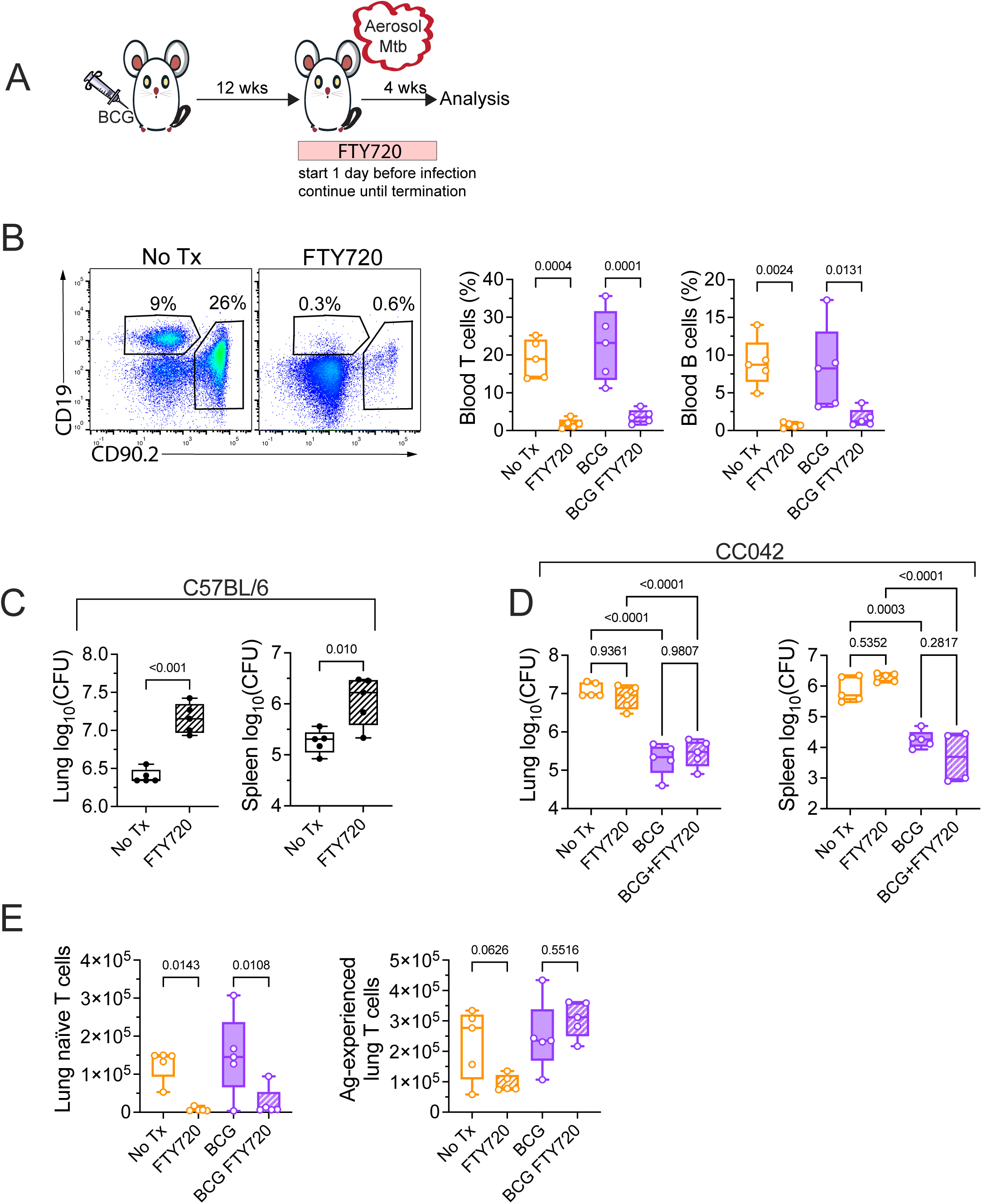
Protection of CC042 mice by BCG is not abrogated by FTY720 treatment. A. Experimental scheme. B6 and CC042 mice were vaccinated with BCG or not and rested for 12 weeks. FTY720 (4 mg/kg) was administered to half of the mice in each group, starting one day before infection and continuing for four weeks. All mice were infected with Rv.YFP and analyzed four wpi. B. Analysis of FTY720 treated CC042 mice. Left, representative flow cytometric analysis of CD19 (B cells) and CD90 (T cells) expression by total live cells in the blood of untreated and FTY720 treated mice. Right, frequency of T and B cells, as a percentage of total live cells, in the blood of different groups of control and vaccinated mice, untreated or treated with FTY720. The data are from one of two independent experiments with similar results, n=5 mice per group. One-way ANOVA with Šídák’s multiple comparisons test. Bars, mean. Error bars, SD. C-D. CFU in the lung (left) and spleen (right) of B6 (C) and CC042 (D) mice at four wpi. (C) Unpaired t-test. Each point represents an individual subject, n=5 mice per group. Bars, mean. Error bars, SD. Data is representative of one of two individual experiments with similar results. (D) One-way ANOVA with Šídák’s multiple comparisons test. Each point represents an individual subject, n=5 mice per group. Bars, mean. Error bars, SD. Data is from one of two independent experiments with similar results. E. Naïve (left, CD44–CD62L+) and antigen-experienced (right, CD44+CD62L–) T cell counts in the lungs of CC042 mice at four wpi. The data is from one of two independent experiments with similar results, n=5 mice per group. One-way ANOVA with Šídák’s multiple comparisons test. Bars, mean. Error bars, SD.

We next enumerated T cells in the lungs of CC042 mice at the 4-week time point. FTY720 treatment reduces the total number of T cells compared to the untreated cohort. In both the unvaccinated controls and the BCG-vaccinated CC042 mice, there is a significant (85-94%) reduction in naïve (CD44^−^CD62L^+^) T cells (Fig.3E, left; and Fig.S3 for gating scheme). In the unvaccinated CC042 mice, FTY720 treatment also significantly reduces the number of antigen-experienced (CD44^+^CD62L^−^) T cells by 71% (Fig.3E, right). However, in BCG-vaccinated CC042 mice, the CD44^+^ lung T cells are not affected by FTY720 treatment, suggesting existence of a population CD44^+^CD62L^−^ T cells elicited by BCG that became lung resident TRM cells prior to Mtb challenge. These data suggest that TRM localization to the lungs can be independent of CD11a and could depend on other integrins such as αE (CD103) for localization. Although the total number of TRM cells is small, these T cells, which were recruited to the lungs of CC042 mice after BCG vaccination might mediate pulmonary protection in CC042 mice.

### The initial phase of BCG-induced protection does not require memory T cells

To determine whether BCG protection of CC042 mice is mediated by TRM cells, we depleted BCG-induced memory T cells by transient administration of anti-CD4/CD8α mAb for three weeks at the end of the rest period (Fig.4A). First, we confirmed that the in vivo depleting antibody doesn’t interfere with the analysis of our target cell populations (Fig.S4A). We established that twice weekly administration of GK1.5 and 2.4.3 mAbs for three weeks before challenge significantly reduced CD4 and CD8 T cells in the lung, spleen and inguinal lymph node (iLN) of CC042 mice (Fig. S4B) and depleted 99% and 94% of circulating CD4 and CD8 T cells in the blood (Fig.4B-C).

**Figure 4.**
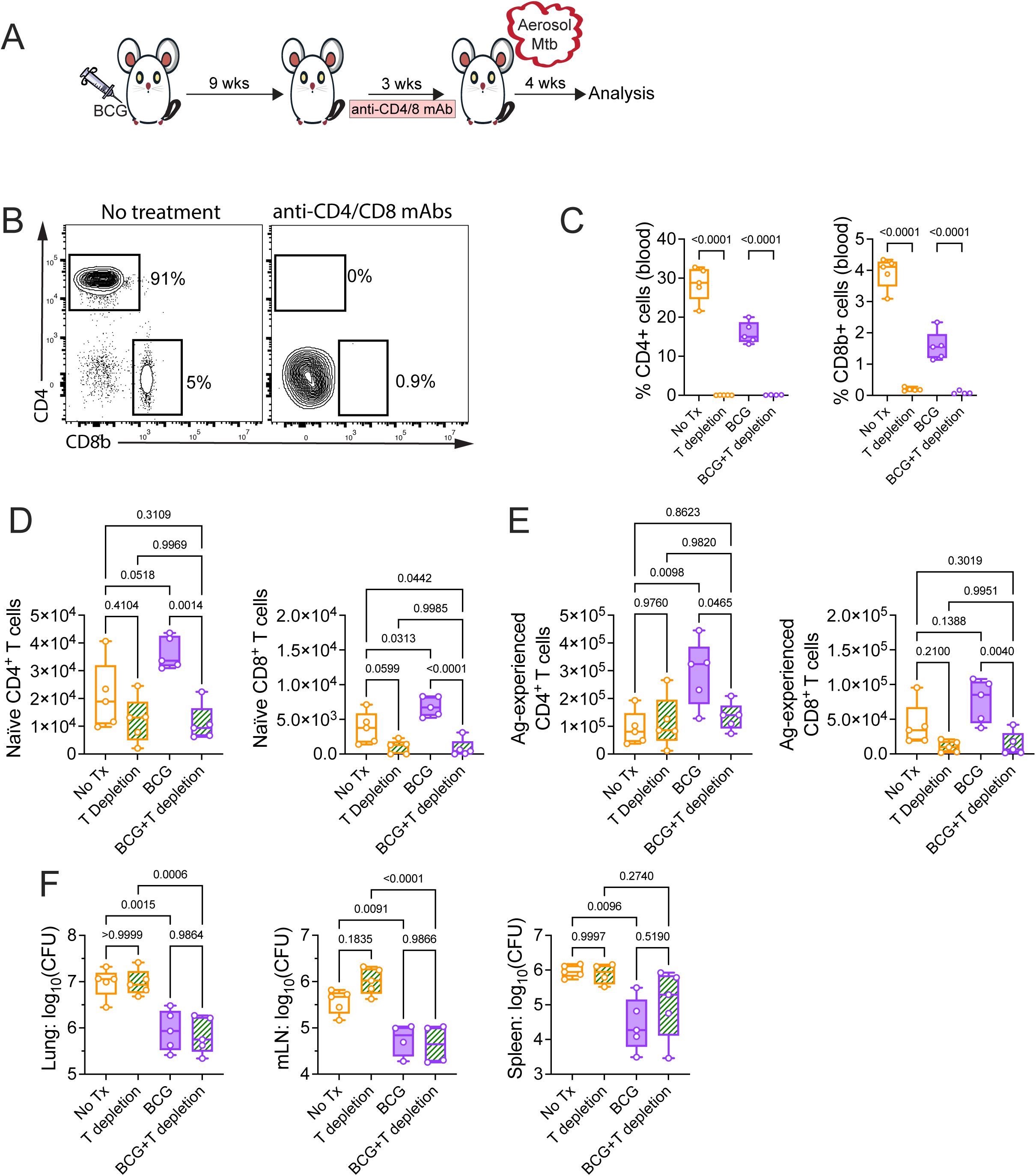
BCG-mediated protection in CC042 mice is independent of CD4 and CD8 memory T cells. A. Experimental scheme. CC042 Mice were BCG vaccinated or not and rested for nine weeks. A cohort from each vaccination group was treated with a combination of anti-CD4 and anti-CD8α mAbs for three weeks. Then, all mice were infected for four weeks with Rv.YFP. B. Representative flow plot of CD4 and CD8β expression of CD90+ T cells in the blood at four wpi of a representative untreated CC042 mouse and one treated with anti-CD4 and anti-CD8α mAbs. C. Frequency of CD90+ CD4 and CD8β T cells in blood at the end of the depletion period before Mtb infection of CC042 mice. Data is from one of two independent experiments with similar results. One-way ANOVA with Šídák’s multiple comparisons test. Each point represents an individual subject, n=5 mice per group. Bars, mean. Error bars, SD. D-E. Enumeration of naïve (CD44–CD62L+, D) and antigen-experienced (CD44+CD62L–, E) CD4 (left) and CD8 (right) T cells in the lungs of CC042 mice four weeks after infection. Results are from one of two independent experiments with similar results, n=5 mice per group. One-way ANOVA with Tukey’s multiple comparisons test. Bars, mean. Error bars, SD. F. BCG-elicited protection in the lung (left), mLN (middle), and spleen (right) is not affected by CD4 and CD8 T cell depletion. One-way ANOVA with Šídák’s multiple comparisons test. Each point represents an individual subject, n=5 mice per group. Bars, mean. Error bars, SD. Data is from one of two independent experiments with similar results.

We counted naïve (CD44^−^CD62L^+^) and antigen-experienced (CD44^+^CD62L^−^) CD4 and CD8 T cells in the lung four weeks after Mtb challenge (Fig.S3 for gating scheme). BCG vaccination boosted the total number of T cells in the lung following Mtb infection as indicated by a modest increase of naïve CD4 and CD8 T cells, and a more significant increase of antigen-experienced CD4 T cells (∼3-fold, p=0.0098; Fig.4D, E). T cell depletion after BCG vaccination prevented the subsequent increase (i.e., boost) of CD4 T cells in the lungs of Mtb challenged mice (Fig.4E). The number of CD44^+^ CD4 T cells in the lungs of the ‘BCG+T depleted’ group was similar to the ‘No Tx’ group, suggesting that depletion of T cells prior to Mtb challenge eliminated memory CD4 T cells primed by BCG that expand in response to Mtb infection.

Last, we enumerated Mtb in lung, mLN and spleen to assess effects of memory T cell depletion on protection. T cell depletion prior to challenge did not adversely affect Mtb burden in unvaccinated mice (No Tx’ vs. ‘T depletion’, Fig.4F). This was not surprising since reconstitution of the CD4 T cell compartment occurs before the endpoint, and T cells have a delayed appearance in the lung of CC042 mice because of their CD11a deficiency (15). Interestingly, BCG-mediated protection was maintained in the lungs, mLNs, and spleens of CC042 mice despite CD4 and CD8 T cell depletion (BCG vs. ‘BCG+T depletion’, Fig.4F). These results suggest that memory CD4 and CD8 T cells are not the major mediators of lung protection seen four weeks after infection in CC042 mice. Instead, these data suggest that early protection conferred by BCG vaccination in CC042 mice is mediated by immune responses that do not require T cell recall responses.

### B cells are not required for BCG-mediated protection of CC042 mice

As BCG stimulated protective immunity despite depletion of memory T cells, we considered whether B cells or antibody responses elicited by BCG could mediate protection against Mtb in CC042 mice. To test this hypothesis, we administered anti-CD20 mAb to deplete B cells. The first dose was given one week before BCG vaccination to prevent initiation of an antibody response to mycobacterial antigens, and because B cell depletion persists for ∼6-8 weeks, a second dose was given five weeks after vaccination (Fig.5A) (25). This strategy was designed to specifically prevent B cell and antibody responses to BCG. We confirmed that the anti-CD20 mAb (MB20-11) used to deplete B cells in vivo did not interfere with the identification of B cell populations using anti-CD19 by flow cytometry (Fig.S4A). This permitted us to verify efficient B cell depletion in CC042 mice (Fig.S5A-B). Anti-CD20 mAb treatment reduces the numbers of B cells in the lung and iLN by 78% and 99%, respectively. Twelve weeks after vaccination, the mice were challenged with low-dose aerosolized Mtb Rv.YFP, and protection was assessed four weeks after challenge. Four weeks after BCG vaccination, circulating B cells in peripheral blood were reduced by 92-94% compared to untreated CC042 mice (Fig.5B). To assess the functional consequences of B cell depletion, antibody titers to culture filtrate protein (CFP) was determined four weeks after Mtb challenge. Although the levels were low and variable, BCG-vaccinated CC042 mice had an increase in their total anti-CFP antibodies compared to untreated mice (Fig.5C). In contrast, B cell depletion prior to vaccination and during the rest period abrogated an increase of antibodies specific for culture filtrate proteins (CFP) (Fig.5C). Antibodies specific for whole cell lysate (WCL) had a similar pattern (Fig.5C). Finally, depletion of B cells before and after BCG vaccination did not disrupt protection of CC042 mice, as the Mtb burden in the lung and spleen of these mice were similarly reduced in the BCG and BCG + B cell depletion groups, compared to untreated mice (Fig.5D). Thus, B cell and antibody responses are dispensable for protection induced by BCG vaccination in CC042 mice.

**Figure 5.**
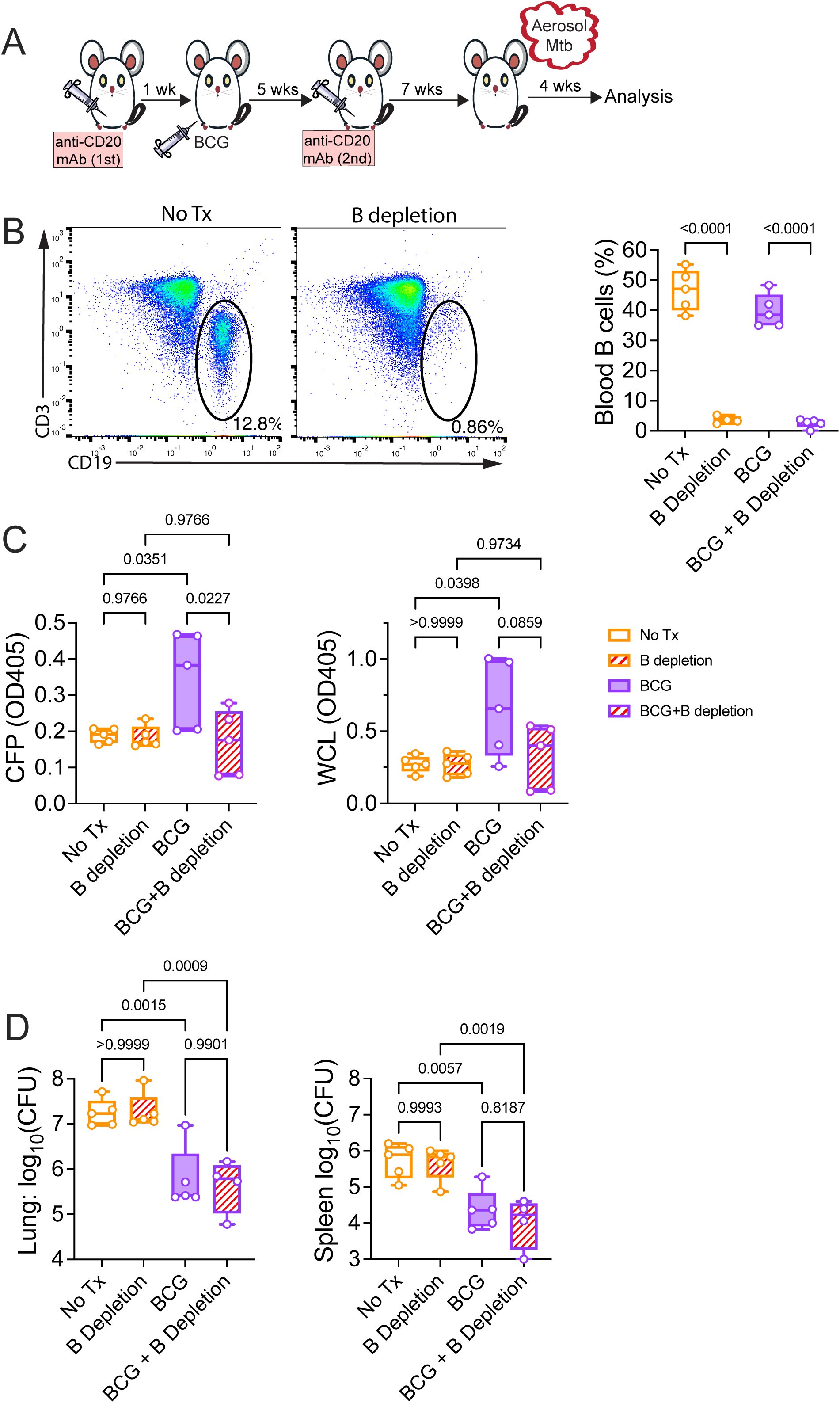
Depletion of B cells does not abrogate BCG-mediated protection in CC042 mice. A. Experimental scheme. CC042 mice were treated with anti-CD20 depletion mAbs one day before and five weeks after BCG vaccination. Mice were rested for 12 weeks after vaccination, and then all mice were infected with Rv.YFP for four weeks. B. Representative flow plots (left) showing CD19 (B cells) and CD3 (T cells) expression of total live cells in the blood of BCG vaccinated versus BCG vaccinated and B cell depleted CC042 mice and frequency (right) of CD19+ B cells in blood four weeks after BCG vaccination of CC042, as determined by flow cytometry. Data is from one of two independent experiments with similar results. One-way ANOVA with Šídák’s multiple comparisons test. Each point represents an individual subject, n=5 mice per group. Bars, mean. Error bars, SD. C. Quantification of total serum antibodies specific to CFP (left) and WCL (right) at the endpoint. One-way ANOVA with Šídák’s multiple comparisons test. Each point represents the mean of technical duplicate replicates, n=5 mice per group. Box plots indicate median (middle line), 25th, 75th percentile (box) and minimum and maximum (whiskers). Data is from one of two independent experiments with similar results. D. CFU in the lung (left) and spleen (right) at four wpi. One-way ANOVA with Šídák’s multiple comparisons test. Each point represents an individual subject, n=4-5 mice per group. Bars, mean. Error bars, SD. Data is from one of two independent experiments with similar results.

### BCG does not disseminate to the bone marrow nor alter hematopoietic stem cells after subcutaneous vaccination

The finding that BCG vaccination of human infants leads to protection from TB and other pathogens led to the idea of cross-immunity and the discovery of innate trained immunity (26–28). In the mouse model, if BCG gains access to the bone marrow (BM) compartment, it remodels hematopoiesis by inducing epigenetic modifications in hematopoietic stem cells (HSCs) (29). The result is that BM-derived macrophages have a greater capacity to restrict intracellular replication of Mtb (29). To assess whether s.c. BCG vaccination of CC042 mice alters the HSC compartment, we quantified BCG in BM eight weeks after subcutaneous vaccination at the hock. No cultivable BCG was detected from the BM of B6 or CC042 mice in two independent experiments, although BCG persisted in other tissues, particularly in the draining LNs of CC042. mice (Fig.6A).

**Figure 6.**
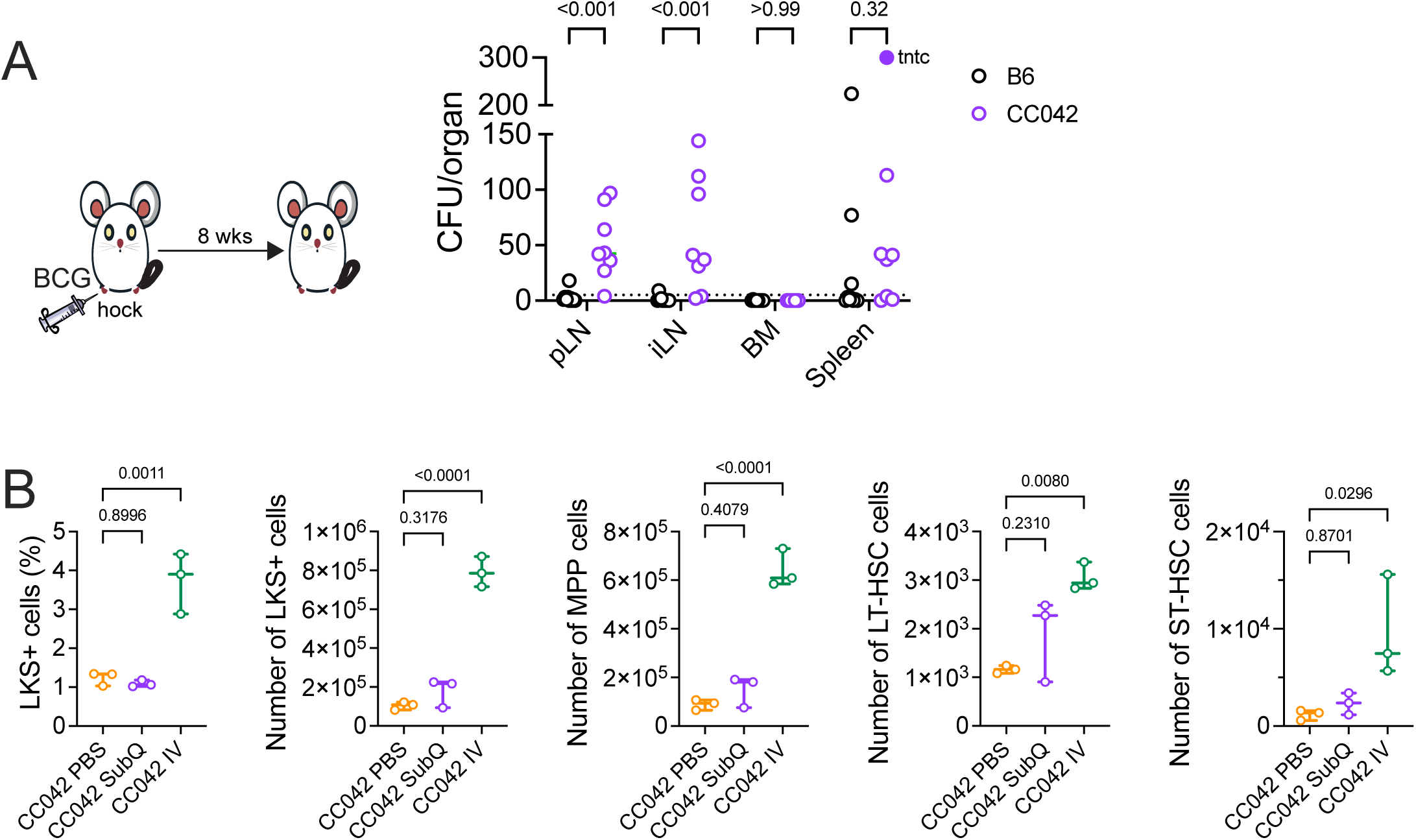
Subcutaneous BCG vaccination of CC042 mice does not result in HSC expansion in the bone marrow. A. B6 and CC042 mice were vaccinated subcutaneously at the right hock and viable BCG recovered eight weeks after vaccination organ is plotted. BCG persists in the ipsilateral (right) pLN, iLN and spleen of both strains. The bone marrow is sterile in both B6 and CC042 eight weeks after vaccination. Data is combined from two independent experiments, n=4-5 mice per group. Mann-Whitney t-test. The limit of detection was five CFU per organ. B. CC042 mice were vaccinated with BCG via the i.v. or s.c. route, or sham-vaccinated with PBS via the i.v. route and the expansion of HSCs was determined four weeks post vaccination in the BM. Representative result of three independent experiments with similar results, n=3 mice per group. One-way ANOVA with Dunnett’s multiple comparisons test. Line, mean. Error bars, SD.

As BCG could transiently reside in the BM or exist below the detection threshold, we next assessed whether there were changes among HSCs following vaccination using two different routes of administration: s.c. and i.v. BCG. The i.v. route was selected to ensure BCG could enter the BM and remodel HSCs (29). As expected, i.v. administration of BCG to CC042 mice significantly altered the HSC in the BM compared to i.v. PBS-treated controls (Fig.6B). CC042 mice vaccinated with i.v. BCG have significant increases in the frequency and absolute number of lineage negative/c-Kit+/Sca-1+ (LKS+) cells, and in the number of multipotent progenitors (MPP), long-term HSC (LT-HSC), and short-term HSC (ST-HSC) compared to the other groups (Fig.6B, and Fig.S6 for gating scheme). Importantly, CC042 mice vaccinated with BCG by our standard protocol (i.e., subcutaneously in the flank), were statistically indistinguishable from non-BCG vaccinated CC042 mice (Fig.6B). As we do not detect any stigma of HSC remodeling, we think it is unlikely that innate training via the expansion of HSCs is the dominant mechanism of protection following s.c. BCG vaccination in CC042 mice.

### T cell responses after BCG vaccination help prime innate-like protection to Mtb

As BCG persists in the draining LNs after subcutaneous vaccination (Fig.6A), we questioned whether persistent BCG could enhance immunity in CC042 mice. To determine whether BCG is present in CC042 mice at the time of Mtb challenge, we determined the burden of BCG in various tissues 12 weeks after s.c. BCG vaccination. Three of ten B6 mice had 5-20 bacilli in their iLN; no BCG was detected in the non-draining LN, spleen or lung (Fig.7A). In contrast, BCG was recovered from the spleen and axillary LN of CC042 mice, and eight of ten mice had bacilli in the iLN, with six having more than 125 CFU (Fig.7A). Given the inability of CC042 mice to clear BCG from the iLN after BCG vaccination, we hypothesized that persistent BCG chronically stimulates anti-mycobacterial immunity and protection of CC042 mice. To test this possibility, cohorts of CC042 mice with were vaccinated with BCG, or left unvaccinated. A third group was vaccinated with BCG and treated with isoniazid and rifampicin in drinking water starting eight weeks after vaccination to clear any persistent BCG. Eight weeks was chosen to permit the immune responses to evolve but ensure sufficient time to kill any persistent BCG. We first determined that following 3.5 weeks of treatment with isoniazid and rifampicin, all viable BCG in the iLN of vaccinated CC042 mice were eliminated, confirming the efficiency of the antibiotic regimen against BCG (Fig.S7A). After treatment for 3.5 weeks, and a four-day washout period for the antibiotic-treated group, (i.e., 12 weeks following BCG vaccination), all mice were infected with low dose aerosolized YFP.Rv for four weeks and protection was assessed by enumerating Mtb in the lung (Fig.7B). Elimination of persistent BCG following BCG vaccination does not result in the loss of protection in CC042 mice as the bacterial burden in the lungs of the BCG-vaccinated and BCG-vaccinated + antibiotic-treated (BCG+abx) cohorts were both significantly reduced compared to the non-vaccinated cohort and not significantly different from each other (Fig.7C). This shows that viable BCG remaining in the iLN of CC042 mice are not required to protect CC042 mice. These data also suggest that BCG induction of protective immunity in CC042 mice occurs within 12 weeks of vaccination.

**Figure 7.**
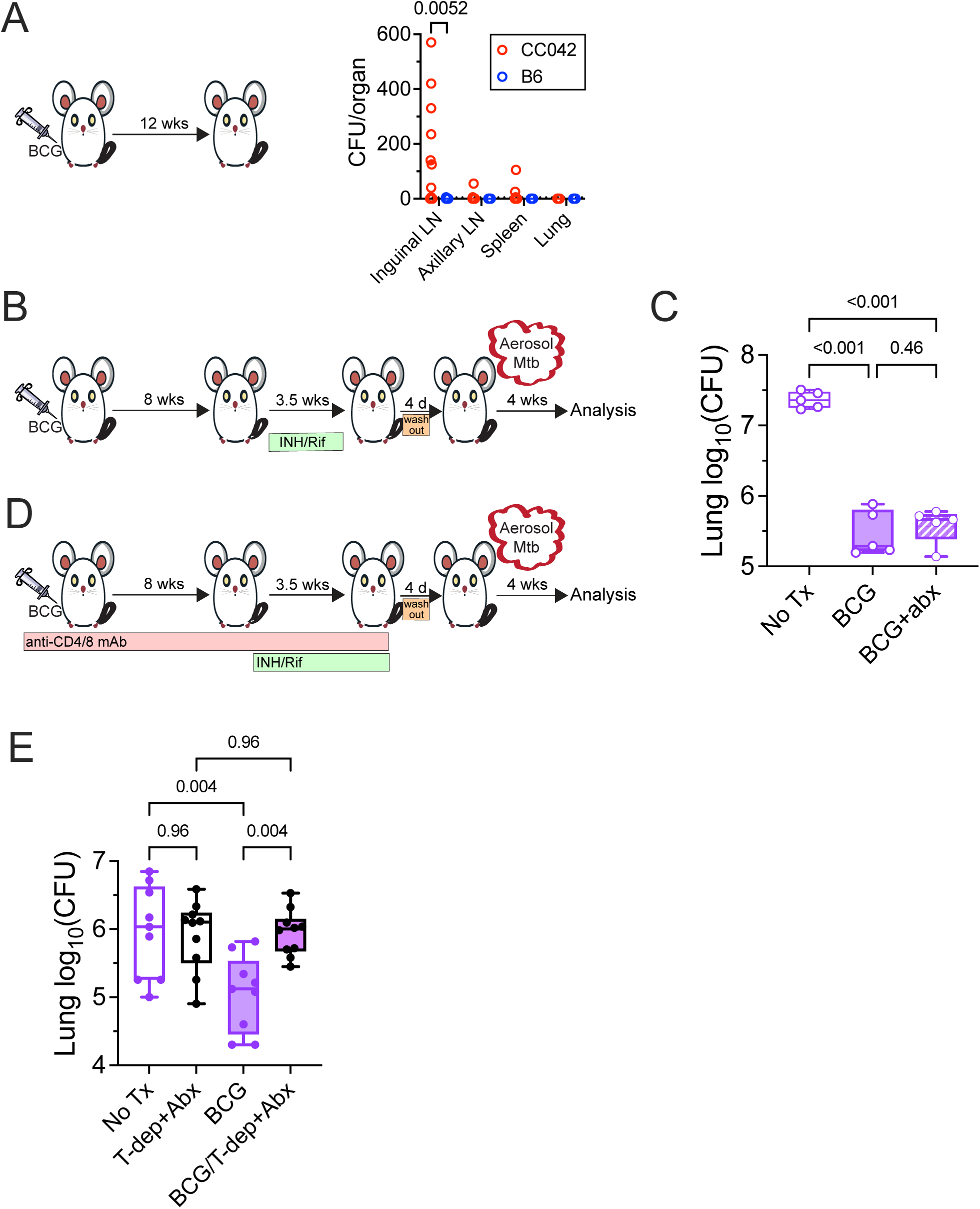
The presence of T cells during vaccination is required for the development of protective immunity. A. B6 and CC042 mice were vaccinated with BCG and 12 weeks later, presence of viable BCG was enumerated in the iLN, axillary LN, spleen and lung. Data is pooled from results of two independent experiments. Each point represents an individual mouse, n=10 mice per group. Mann-Whitney t-test. The limit of detection (LOD) was five CFU. B. Experimental scheme. CC042 mice were vaccinated with s.c. BCG or not and then rested for eight weeks. Half of the BCG-vaccinated mice were then provided drinking water with both isoniazid and rifampicin for 3.5 weeks. 12 weeks after BCG vaccination, all mice were infected for with Rv.YFP for four weeks. C. CFU in the lung. Data is representative of two independent experiments with similar results. One-way ANOVA with Šídák’s multiple comparisons test. Each point represents an individual subject, n=4-5 mice per group. Bars, mean. Error bars, SD. D. Experimental scheme. CC042 mice were treated with CD4 and CD8α depletion antibodies starting one week before BCG vaccination and for the duration of the 12 week rest period. Nine weeks after vaccination, all mice within the depletion group were provided drinking water with both isoniazid and rifampicin for 2.5 weeks. 12 weeks after BCG vaccination, all mice were infected with Rv.YFP for four weeks. E. CFU in the lung. Data is from one of two independent experiments with similar results. One-way ANOVA with Šídák’s multiple comparisons test. Each point represents an individual subject, n=5 mice per group. Bars, mean. Error bars, SD.

While memory CD4 and CD8 T cells do not mediate the effector phase of BCG-elicited protection against Mtb infection (Fig.4), we asked if the protective immune response elicited by BCG in CC042 mice requires T cells early during the immune response. To answer this question, we depleted CD4 and CD8 T cells commencing one week before BCG vaccination and continued mAb administration through the duration of the vaccine rest period (Fig.7D). Since CC042 mice are unable to clear BCG as efficiently as B6 mice, and we expected BCG to replicate more in the absence of T cells, we prevented BCG persistence by treating the mice with isoniazid and rifampicin as described above. Following a four-day washout period, the mice were infected with low-dose aerosolized Mtb YFP.Rv, and after four weeks, protection was assessed (Fig.7D). Here, sustained depletion of CD4 and CD8 T cell populations in BCG vaccinated CC042 mice resulted in 99% depletion of both CD4 and CD8 T cells in the peripheral blood as determined by flow cytometry (Fig. S7B). Continual depletion of CD4 and CD8 T cell cells prior to BCG vaccination and during the rest period, eliminated BCG-mediated protection as the bacterial burden in the lung (and spleen) in this group was the same as the untreated group (“No Tx”, Fig.7E). Thus, while the enhanced capacity of BCG-vaccinated mice to restrict Mtb replication in the lung is not mediated by memory CD4 and CD8 T cells, CD4 and CD8 T cells are nonetheless required during the response to BCG to prime innate-like protection against Mtb challenge.

### Long-term BCG-mediated protection of CC042 mice against Mtb infection requires T cells

BCG vaccination generates protective immunity, as defined by lung CFU reduction at the standard time point of four weeks after Mtb challenge in CC042 mice, which is dependent on CD4 and CD8 T cells, but not mediated by memory T cell responses (Fig. 1, 4, 7). Having shown that BCG vaccination of CC042 mice extends survival, a measure of the durability of immunity (Fig.1), we considered whether protection at the early and late phases of infection were mediated by different immune mechanisms. We hypothesized that innate mechanisms of protection activated by BCG reduce the lung Mtb burden four weeks after challenge, while T cells mediate long-term protection. To test the latter possibility, CC042 mice were vaccinated with BCG, T cells depleted after BCG vaccination during the rest period, and following Mtb challenge, survival was determined using weight loss as a humane endpoint (Fig.8A). We used flow cytometry to determine that the total T, CD4 and CD8 T cell populations of the BCG-vaccinated CC042 mice were 77%, 96%, and 99% depleted, respectively, in the peripheral blood (Fig. S7C).

**Figure 8.**
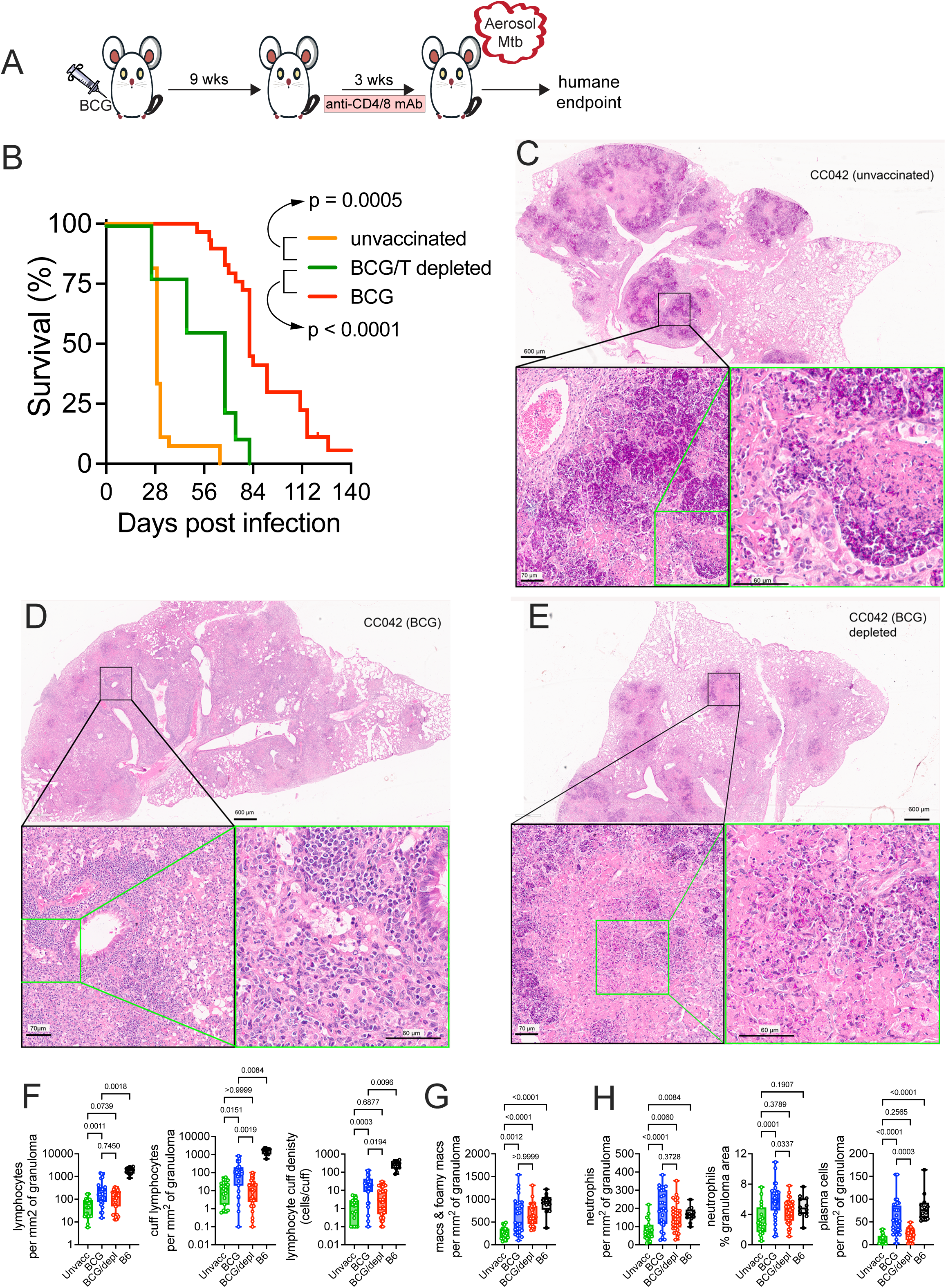
BCG-induced long-term protection of CC042 mice against Mtb requires T cells. A. Experimental scheme. CC042 mice were BCG vaccinated or not and rested for nine weeks. After nine weeks, half of the BCG vaccinated mice were treated with a combination of CD4 and CD8α depleting antibodies for three weeks. Then, all mice were infected for four weeks with Mtb Erdman. B-D. Unvaccinated (n=27), BCG vaccinated (n=30) and BCG-vaccinated plus CD4 + CD8α depleted (n=9) were monitored until they reached a predetermined humane endpoint and were euthanized. Data is pooled from three experiments. The difference between non-vaccinated and BCG-vaccinated CC042 mice reaches statistical based on the Mantel-Cox log-rank test (p < 0.0001). The difference between non-vaccinated and BCG-vaccinated and treated CC042 mice reaches statistical based on the Mantel-Cox log-rank test (p = 0.0005). Representative images of lung pathology at the time of death from an unvaccinated CC042 mouse (B), BCG-vaccinated CC042 mouse (C) and BCG-vaccinated and T cell depleted CC042 mouse (D). Green and black boxes denote areas of higher magnification within the same image.. F-G. Automated image analysis of histopathological tissue. Three to five lung lobes were analyzed per mouse for a total of 19 (unvaccinated), 27 (BCG), 34 (BCG+T depletion), and 12 (B6) lobes. To account for variability in the tissue sections, the analysis is normalized to the granuloma area. Comparisons between different groups were analyzed using a non-parametric one-way Anova (Kruskal-Wallis). **F**. The total number of lymphocytes in the lesions (left), number of lymphocytes in cuff regions (middles), and the density of lymphocytes per cuff (right). **G**. The total number of granuloma macrophages. **H**. The total number of neutrophils in the lesions (left), the percent of the granuloma area occupied by neutrophil infiltrates (middles), and the number of plasma cells.

As previously observed (Fig.1I), non-vaccinated CC042 mice rapidly succumb to pulmonary TB, and BCG vaccination significantly extends survival. When CD4 and CD8 T cells are depleted after BCG vaccination, but before Mtb infection, the median survival is significantly reduced from 82 days to 68 days (p<0.0001, Fig.8B), which indicates that memory T cells elicited by BCG are important in mediating long term protection of CC042 mice conferred by vaccination. Conversely, despite memory T cell depletion before Mtb challenge, the median survival is significantly increased from 28 days to 68 days, suggesting that an unknown component of long-term protection by BCG is independent of vaccine-induced T cell recall responses (p=0.0005, Fig.8B).

To understand how memory T cells elicited by BCG modify host immunity and TB pathogenesis, we collected lung tissue from mice that had reached humane endpoint at the time of euthanasia. Fixed tissue sections stained as before (Fig. 2) with an AFB plus H&E stain were evaluated by a veterinary pathologist (Fig.8C-E). In response to Mtb infection, unvaccinated CC042 mice developed necrotizing inflammation characterized by a paucity of lymphocytes and plasma cells with marked fibrin exudation into alveolar spaces, pyknotic nuclear debris (consistent with neutrophil and macrophage necrosis) that obstructed bronchioles, and massive granuloma necrosis (Fig.8C). Numerous extracellular AFB were primarily located extracellularly in the inflammatory exudate. These histopathological findings suggest that Mtb infection and the necrotic response to bacilli in unvaccinated CC042 mice damages capillary endothelial cells and alveolar septae, due to bacterial molecules, inflammatory cytokines, alarmins, or other products from necrotic cells.

BCG vaccination of CC042 mice changed the nature of the infiltrates from a severe acute fibrinonecrotizing response without lymphocytes to lesions containing fewer and smaller fibrinonecrotic foci surrounded by foamy macrophages and edema, and many prominent perivascular and peribronchiolar cuffs containing numerous lymphocytes and plasma cells (Fig. 8D). AFB were detected in both intracellular (macrophages) and extracellular (fibrin, necrotic) locations. These findings, in conjunction with survival extension, suggest that main protective effects of BCG vaccination in the lungs are (i) reduced cellular necrosis and fibrin exudation; (ii) increased numbers of lymphocytes, plasma cells, and macrophages; and (iii) restriction of bacilli growth to intracellular compartments, likely by improved macrophage activation.

The lung histology from BCG vaccinated, memory-T cell depleted mice was more variable. Some features correlated to survival, for example, lung lesions in mice that succumbed early resembled unvaccinated CC042 mice, characterized by fibrinonecrotizing inflammatory foci with necrotic neutrophils and/or macrophages occupying approximately 40-60% of the lung area, with numerous extracellular AFBs and a paucity of lymphocytes and plasma cells (Fig 8E). In contrast, the lungs from memory-T cell depleted mice that survived longer were infiltrated approximately 80% of the lung tissue section by non-necrotic macrophages and foamy macrophages containing fewer AFBs (primarily intracellular single bacteria, and few clusters of bacilli) and sparse lymphoid aggregates. The lungs of survivors did not contain abundant fibrin exudation into alveolar spaces or contain regions of alveolar septal necrosis.

To quantify lung granuloma regions and cell types of interest, we applied a machine learning model that was trained and validated using the Aiforia platform. Total lung tissue section area, granuloma area, and percent of lung tissue section occupied by granuloma was similar across the groups (Fig.S8A). The regions of cell-poor necrosis and accumulation of pyknotic debris were ameliorated by BCG vaccination, irrespective of memory T cells depletion (Fig.S8B). BCG-vaccinated CC042 mice had more ∼4-fold more lymphocytes in lung granulomas than unvaccinated mice; however, B6 mice had 10-fold more lymphocytes than BCG-vaccinated CC042 mice (Fig.8F). We find that lymphocyte cuffs are associated with protection. Lymphocyte cuff formation is stimulated by BCG vaccination but is independent of memory T cells. BCG vaccinated mice have ∼10-fold more lymphocytes in the cuff regions than unvaccinated mice, and the increased number of lymphocytes is abrogated when memory T cells are depleted before Mtb challenge (Fig.8F). Thus, the lymphocyte density in each cuff correlates with protection (∼13-fold greater in vaccinated mice) and dependent upon anamnestic T cell responses (Fig.8F). Unexpectedly, plasma cells similarly correlated with protection and were T cell dependent (Fig.8H). Macrophages (Fig.8G) and neutrophils (Fig.8H) were significantly increased in the lesions of BCG vaccinated mice and their numbers were unaffected by memory T cell depletion.

Enumerating bacilli in tissue sections is a challenge because Mtb do not grow as single bacilli and are not distributed uniformly. Our model identifies three forms of Mtb: single bacterium, clusters, and biofilms. The single AFB and clusters are localized in macrophages or in debris (i.e., extracellular). Interestingly, the effect of vaccination was two-fold. It led to a reduction in extracellular AFB clusters and AFB biofilm (Fig.8C, D), and to an increase in the number of intracellular single AFB (Fig.S8E). Both effects appear to be independent of memory T cells. Thus, BCG vaccination of CC042 mice alters the long-term host response to Mtb that can be seen at the time of death. It increases macrophage recruitment, reduces necrotic damage, and shifts that location of the Mtb bacilli from an extracellular location to an intracellular location. These effects appear to be independent of conventional memory CD4 and CD8 T cells. Additionally, there is an increase in lymphocyte cuff formation and lymphocyte density, plasma cells, and neutrophils, all of which does appear to be dependent on the presence of memory T cells at the time of Mtb challenge.

## Discussion

A defective *Itgal* gene, which encodes the α-subunit of the integrin molecule CD11a, increases the susceptibility of CC042 mice to Mtb infection (15). A major role of CD11a is lymphocyte trafficking to secondary lymphoid organs and sites of infection. Specifically, CD11a mediates cell extravasation across endothelium (16). Just as important is its involvement in immune synapse formation. CD11a facilitates firm, stable connections between antigen presenting cells and T cells, promotes immune synapse formation, establishes signaling cascades and potentiates the release of cytotoxic granules (17). Thus, the pervasive lung and granuloma necrosis, and early death following Mtb infection are all consistent with the inability of CC042 mice to produce functional CD11a protein (15). Surprisingly a screen of CC mouse strains showed that BCG vaccination of CC042 mice leads to a significant reduction of lung CFU four weeks after infection (10). Here, we confirmed that s.c. BCG vaccination of CC042 mice leads to significant reductions in Mtb burdens in the lung, spleen and lung-draining mLN, compared to unvaccinated control mice, measured four weeks after low dose aerosol Mtb infection, and increases their survival 3-fold. The reduction in lung CFU and increased survival are both standard parameters used to assess vaccine-induced protection in the murine TB model. Therefore, we anticipated that the study of mouse strain CC042 would reveal non-conventional mechanisms of immunity elicited by BCG that mediate protection against Mtb infection.

We were surprised to detect an increase in lymphocytes in the lung lesions of BCG vaccinated CC042 mice after Mtb challenge, and many of these are T cells based on flow cytometric analyses. Together with the persistence of T cells in the lung despite FTY720-treatment, this suggests that their trafficking to the lung could be dependent upon adhesion molecules other than LFA-1 (e.g., αEβ7), and they might have been derived from lung TRM. However, a CD4 and CD8α depletion strategy used after vaccination, which depleted antigen-experienced T cells, including those in the lung, and permitted reconstitution of T cell immunity to Mtb, did not affect the ability of BCG to reduce Mtb abundance in the lungs of CC042 mice. Similarly, depletion of B cells before vaccination, a strategy designed to prevent antibody responses to BCG, did not affect protection. Although BCG-elicited protection is independent of CD4 and CD8 T cells and B cells, as defined by the reduction in Mtb CFU in the lung four weeks after infection, sustained protection requires T cells. Thus, using the same T cell depletion strategy, we show that the survival benefit conferred by BCG to CC042 mice is disrupted when BCG-induced T cells are depleted. Thus, we observe two distinct effector phases of immunity in CC042 mice: an initial phase that does not require the presence of CD4 and CD8 T cells, and a later T cell dependent phase.

There is still considerable uncertainty about how BCG protects against infection. Some theories posit that BCG primes adaptive immune cells, that are poised for rapid and efficient recall responses upon Mtb infection. BCG-primed T cells transfer protection to immunodeficient mice, although it is uncertain which T cell subsets contribute to long lasting immunity. While it has been relatively straightforward to show that CD4 T cells correlate with protection, increasingly, new technical approaches show a correlation between CD8 T cells and immunity (30, 31). Other T cell subsets are generated during BCG vaccination, but their role in protection is still being sorted out (32, 33). While evidence is against a role for B cells and antibody responses as important in the mouse TB model most of the data was generated using the C57BL/6 inbred strain of mice, which has a strong Th1 responses to mycobacteria including *M.tb* (34) and that Th1-biased phenotype may mask protective roles for B cells in the C57BL/6 strain. Recently, data is accumulating that B cells may have a more important role, in other mouse strains (35) humans and NHPs, particularly in the context of anamnestic responses (36).

Other theories suggest that BCG remodels innate immune responses to become more potent responders to a broad range of pathogens, including Mtb (29, 37). The latter idea arose from data that BCG vaccination at birth protects infants from Mtb and other pathogens, leading to a reduction in all-cause mortality compared to unvaccinated infants (28). The term given to this phenomenon is trained innate immunity. BCG and other stimulants of innate immunity (e.g., yeast), epigenetically modify hemopoietic stem cells (HSC), especially of the myeloid lineage (38, 39). However, whether innate immune training by BCG vaccination contributes to protection in the mouse TB model is unclear. Vaccination of mice with BCG by the intravenous route, but not the s.c. or intranasal route, induces “training” and BM-derived macrophages have an enhanced ability to restrict Mtb growth in vitro (29). The BCG route-specificity appears to reflect a requirement that BCG gains access to the BM compartment. Innate immune training is IFNγ- dependent, and could require T cells, although there are other sources of IFNγ (e.g., natural killer cells). So, while BCG can induce innate immune training, it is not yet clear whether BCG consistently disseminates to the BM in mice, non-human primate (NHP), or humans following s.c. vaccination. Besides innate training (i.e., epigenetic modification of HSCs) is not the only mechanism by which BCG can modify innate responses as it also modifies AM responses (40). It is not yet clear whether innate training leads to durable protection or whether it could be a major mechanism of protection in vivo.

Although animal models do not fully recapitulate human disease, they are indispensable for pre-clinical vaccine development and assessment as they allow for standardized testing of varied conditions and evaluation of different end points and mechanisms. In the early 2000’s, the NIH and the FDA urged harmonization of the work being done in different labs around the world by advocating for using a consistent Mtb challenge strain (i.e., low-dose aerosolized Mtb Erdman) and analysis at a standard 30 days post infection timepoint (41). Increasingly, investigators have questioned how to interpret these early timepoints, as vaccine-induced protection, particularly after BCG vaccination, is often transient. Similarly, the dominant use of C57BL/6 and BALB/c mice in vaccine studies risks optimizing immunization strategies for a single genetic background and genetically controlled immune mechanisms. An ongoing debate is whether to use susceptible mouse strains (42, 43). Previously, there was resistance to the use of susceptible strains such as C3H substrains because of concern that they had defective immunity; however, their development of necrotic and hypoxic lung lesions is more like human granulomas (44, 45).

Surprisingly, despite the absence of CD11a, CC042 mice generate a complex immune response to BCG and allowed evaluation of several disease and immunological parameters and following BCG vaccination and Mtb challenge. Protection induced by BCG vaccination of CC042 mice is based on three parameters – reduced bacillary load, better Mtb containment and less lung disease, and increased survival. Finding that CD4 and CD8 T cells are dispensable for BCG-induced protection four weeks after Mtb challenge is surprising as this is a standard time point to assess vaccine-induced protection in the mouse TB model (45). As we are depleting T cells after BCG vaccination but before Mtb challenge and providing time for reconstitution of the T cell compartment, we are primarily removing memory T cells from the immune repertoire and preventing T cell recall responses (i.e., secondary responses) from developing following Mtb challenge. Consequently, we infer that this early phase of protection is primarily mediated by non-T cell mechanisms of immunity. Although CD4 and CD8 T cells are not acting as effectors during this phase of infection, it appears that T cells are orchestrating the protective immune responses. Depletion of CD4 and CD8 T cells before vaccination prevents the development of protective immunity, which implies that T cells enhance innate responses to control Mtb early during infection (i.e., 4-wpi). While we have not formally identified the immune mechanisms that operate during this phase, it is likely to be an innate immune mechanism (e.g., innate training, natural killer (NK) cells, or donor-unrestricted T cell responses). In contrast, long-term protection (i.e., survival) requires T cells. An important future question is whether these different parameters are interconnected or a reflection of different disease processes and mechanisms of immunity. A focus of many vaccine studies is the identification of immune correlates of protection. Reliance on a single early timepoint would have limited our analysis and failed to uncover a role for T cells in BCG-mediated protection of CC042 mice.

CC042 mice are a complex mosaic of eight highly diverse parental founders. While a defective *Itgal* gene is an important driver of their susceptibility to *Mtb* and *Salmonella* (15, 46), the 15-base intronic deletion is a private mutation that is present in CC042 mice but not any of the founders. There are other genetic loci that affect the susceptibility of CC042 mice to Mtb infection other than the *Itgal* gene. Indeed, CC042 mice are more susceptible than CD11a^−/–^ mice (backcrossed to B6 mice, and the nature of their lung lesions differ significantly (19). Nevertheless, these data show that CD11a is not necessary for BCG-mediated protection to Mtb infection, and immune mechanisms other than conventional B and T cells can be sufficient to control Mtb infection. Based on these studies, we expect that interrogating how genetic diversity affects immunity should provide insight into host resistance against Mtb infection.

## Methods

### Sex as a biological variable

Our study examined male and female animals. The susceptibility of CC042 mice to Mtb infection and the protection elicited by BCG was not affected by sex (see Fig.1). Therefore, sex was not considered as a variable in this study, although both mice of both sexes were used in the experiments and had similar results.

### Statistical analysis

Statistical analyses were performed using GraphPad Prism (v10) software. P values were calculated using unpaired or paired t test, one-way ANOVA, or two-way ANOVA with post-tests as indicated in the figure legends.

### Study approval

Studies were conducted using the relevant guidelines and regulations and approved by the Institutional Animal Care and Use Committee at the University of Massachusetts Chan Medical School (Animal Welfare A3306-01), using the recommendations from the Guide for the Care and Use of Laboratory Animals of the National Institutes of Health and the Office of Laboratory Animal Welfare.

### Mice

All *in vivo* experiments were performed with sex- and age-matched mice, born within three weeks of each other. 6–8-week-old B6 mice were purchased from Jackson Laboratories. CD11a^−/–^ mice were purchased from Jackson Laboratories and bred in house. CC042 (CC042/GeniUnc) mice were initially obtained from the UNC Systems Genetics Core Facility and bred in house. All animals in the study were conducted in the Animal Medicine Facility at the University of Massachusetts Chan Medical School.

### In vivo infections

Frozen Mtb stocks of Rv.YFP (H37Rv expressing yellow fluorescent protein sfYFP) (47) or Erdman were thawed and sonicated for 1 minute and then diluted into 5 mL of CFU buffer (0.01% Tween-80 (Fisher) in PBS (Gibco). Then, mice were exposed to an aerosol generated by the Glas-Col Inhalation Exposure system using bacterial suspension. The average number of bacteria delivered to each mouse was determined by lung CFU assay (n=4-5 mice) within 24 hours after infection and ranged from 20-119 (Mtb Rv.YFP) and 17-100 CFU (Erdman) CFU/mouse.

### BCG vaccination

Frozen stocks of BCG-SSI were thawed on ice for 30 minutes. BCG was triturated through a 22-gauge needle (BD Biosciences) on a 1 mL syringe (BD Biosciences) at least 10 times to disrupt clumps. The bacterial suspension was diluted in PBS to administer 500,000 CFU of BCG in 100 µL (flank vaccinations) or in 20 µL (hock vaccinations).

### Survival Analysis

Infected mice were monitored at least once weekly using the Body Condition Score (BSC) and weight monitoring. Mice with a BSC score ≦2 or weighed <80% of their starting weight were humanely euthanized.

### In vivo depletions

CD4 and CD8α: Mice were given both anti-CD4 (GK1.5) and anti-CD8α (2.4.3) mAbs (Bio X Cell) via the intraperitoneal route twice weekly for the duration of the depletion period. The first dose contained 200 µg of each mAb suspended in 200 µL of PBS, and all subsequent doses contained 100 µg of each mAb suspended in 200 µL of PBS.

CD20: Mice were given 250 µg of anti-CD20 mAb (MB20-11, Bio X Cell) suspended in 200 µL of PBS via the intraperitoneal route one week before vaccination and again six weeks later (five weeks post-vaccination).

Crossblocking analysis: Splenocytes from CC042 mice were incubated individually with 100 µg each of antibody (anti-CD20, MB20-11; anti-CD4, GK1.5; or anti-CD8α, 2.4.3) for 20 minutes at RT. The cells were then washed once with PBS and stained with fluorescently labelled mAbs specific for CD20, CD4, or CD8, as indicated in Figure S4A, and then analyzed by flow cytometry.

### Bacterial burden determination

Vaccinated and infected mice were euthanized at the indicated time points. The left lung lobe, mediastinal lymph node, inguinal lymph node or spleen were homogenized with 2 mm zirconium oxide beads (Next Advance) in a FastPrep homogenizer (MP Biomedical), three rounds for 20 seconds each. Homogenates were then serially diluted and plated on 7H11 plates (Hardy Diagnostics), and CFU was enumerated 21-25 days after incubation at 37C.

### Cell preparation

Lung single cell suspensions were prepared first by homogenizing in a GentleMACS tissue dissociator (Miltenyi) followed by a 30-minute incubation in collagenase (300 units/mL (Sigma) in 6 mL of total RPMI (Gibco). Then, lungs were homogenized for a second time in the GentleMACS before being sequentially filtered through 70 µM and 40 µM cell strainers (Fisher Scientific) and finally suspended in complete RPMI (10% FBS (Gibco), 2 mM L-glutamine (Gibco), 50,000 units penicillin/streptomycin (Gibco), 1 mM sodium-pyruvate (Gibco), 1x non-essential amino acids (Gibco), 1x essential amino acids (Gibco), 25 mM of HEPES (Gibco), 7.5 mM of sodium hydroxide and 0.55 µM 2-mercaptoethanol (Gibco). Spleen and LN single cell suspension were prepared by manual homogenization with plungers from 5 mL syringes (Fisher Scientific) and then were sequentially filtered through 70 µM and 40 µm cell strainers and finally suspended in complete RPMI. Lung and spleen cells were treated with ACK lysis buffer (Gibco) for 1-2 minutes to lyse red blood cells before filtering through 40 µM strainer. BM cells were acquired by flushing the tibia and femur of mice with PBS and then filtering the cells through a 70 µM cell strainer. Cells from single cell suspension were counted on a TC20 (Bio-rad) at 1:20 dilution in PBS.

### Blood lymphocyte preparation

Lymphocytes from blood via cheek bleeds were collected in 2 mL of RPMI containing 80 units of heparin (Fisher Scientific). Then, 1mL of Lympholyte (Cedarlane) was layered beneath the blood and centrifuged for 20 minutes at 1800 rpm. The lymphocyte layer was transferred to a fresh tube containing 2 mL of autoMACS (Miltenyi) running buffer and washed once with PBS before beginning the flow cytometry staining process.

### Serum collection

Blood was collected from mice at the time of sacrifice from the vena cava. The blood was incubated at 4C overnight to form a clot. Then, the blood was centrifuged at 2000g for 10 minutes at 4C. The serum was collected, then filtered through a Multiscreen 96-well 0.22 µM filter plate (Millipore) and frozen at −80C until use.

### Antibody ELISA

Polystyrene 96-well plates were coated with 10 µg WCL or CFP (BEI resources) in PBS overnight at 4C. The plate was blocked with 100 µL per well of 1% bovine serum albumin (Fisher Scientific) in PBS. Then, 50 µL of the serum diluted 1:40 in 1% bovine serum albumin (BSA) was added per well. Next, 50 µl of alkaline phosphatase-conjugated goat anti-mouse IgG (H+L) (Jackson ImmunoResearch) at 1:1000 diluted in 1% BSA was added to each well. Lastly, 50 µL of phosphatase substrate (Millipore Sigma) was added to each well and incubated until the reaction completed, within 30 minutes. The reaction was quenched with 1 M sodium hydroxide (Spectrum Chemical) in water and at OD_405_ on plate reader. All incubations were performed at RT shaking gently and the plate was washed three times with wash buffer (PBS + 0.1% Tween-20 (Fisher Scientific) in between each step.

### Flow cytometry

Cells were washed once with PBS and then stained with Zombie viability dye (Biolegend) for 10 minutes at RT. Cells were then washed with autoMACS running buffer and incubated with 1% anti-mouse CD16/32 (Biolegend) suspended in autoMACS running buffer for 10 minutes at RT. Surface staining then followed by incubating cells with antibodies suspended in autoMACS running buffer for 20 minutes at 4C. Cells were then fixed in 1% PFA (Thermoscientific) suspended in PBS for at least 20 minutes. Samples were acquired on the Miltenyi MACSQuant 16 or the Cytek Aurora. All data was analyzed with FlowJo software, version 10.

### Antibiotic Treatment

Mice were provided drinking water containing 0.5 g/L of isoniazid (Sigma, from 250x stock in water) and 0.1 g/L of rifampicin (Sigma, from 250x stock in DMSO (Fisher Scientific) in the drinking water ad lib at the specified timepoints. Medicated water was protected from light and replaced once a week.

### Histopathological tissue preparation

Lungs were inflated and immersion-fixed immediately after euthanasia with Z-fix (Anatech) for at least 30 minutes and then washed with PBS. Lung tissue was then embedded in paraffin, sectioned, affixed to glass slides and dewaxed in an oven at the Morphology Core Facility at UMass Chan Medical School. Fixed tissue was deparaffinized in a solution of 1 part peanut oil (Planters) to 2 parts xylene (Fisher Scientific) twice for 12 minutes each. Then, the tissue was stained with carbol fuchsin (5% phenol, 10% ethanol, 1% basic fuchsin in distilled water) for 25 minutes, differentiated in acid alcohol (1% hydrochloric acid in 70% ethanol), stained with Harris hematoxylin with glacial acetic acid (Polyscientific) for three minutes, neutralized in bluing reagent (0.25% ammonium hydroxide in distilled water) and counterstained with eosin phloxine alcoholic working solution (Polyscientific). Lastly, the slides were dehydrated in ethanol (Fisher biologics), then xylene and cover slipped with permount (Fisher biologics).

### Histopathological tissue evaluation

Stained lung tissue sections on glass slides were digitally scanned by Aperio ScanScope or AT2 scanners at 0.23 microns/pixel at Vanderbilt University Medical Center’s Digital Histology Shared Resource (Nashville, TN, USA). The digital images were uploaded to the Aiforia Create (v.6.0) platform (Aiforia Technologies, Helsinki, Finland) Slide Viewer for blinded qualitative evaluation and semi-quantitative assessment of tissue architecture, cell types, and AFBs by a board-certified veterinary pathologist (GB).

### Automated image analysis of histopathological tissue

Manual training annotations were provided using the Aiforia Create (v.6.0) platform with default parameter settings (Aiforia Technologies, Helsinki, Finland) to create a multi-layer, multi-class convolutional neural network-based model for automated image analysis to detect and quantify lung tissue, granulomas, subregions of granulomas, i.e., pyknotic debris, cell-poor necrosis, lymphocytic cuffs, fibrin, and viable regions of granulomas containing foci of immune cells, e.g., lymphocytes, macrophages, neutrophils, and plasma cells and their nuclei; and AFB single bacilli, clusters of bacilli, and regions where bacilli formed a mat resembling a biofilm containing too-numerous-to-count organisms. Many iterative rounds of training, testing, and validation by pathologists’ inspection of model performance on independent images (not used in training) were performed until errors were minimized.

## Data availability

All other data are available in the main text or the supplementary materials. Any additional information required to reanalyze the data reported in this paper is available from the lead contact upon request. Further information and requests for resources and reagents should be directed to and will be fulfilled by the lead contact, Samuel M. Behar.

## Acknowledgements

We thank the UMass Chan Flow Cytometry Core for their expertise. We also thank Beshair Nurhussien, Tasfia Rakib, Ellen Acheampong, Seden Bedir and Anthony Tran for technical assistance when performing experiments. Whole slide imaging was performed in the Digital Histology Shared Resource at Vanderbilt University Medical Center (vumc.org/dhsr). This project has been funded by P01 AI123286 (SMB), R01 AI172905 (SMB, GB), R01 HL145411 (GB), and Contract No. 75N93019C00071 (SMB) from the National Institute of Allergy and Infectious Diseases, National Institutes of Health, Department of Health and Human Services. AFO received support from The Immunology and Microbiology Training Grant T32AI007349 and The Medical Scientist Training Program Grant T32GM159591.

## Author Contributions

Conceptualization, AFO and SMB; Investigation, AFO, RL and KC; Formal analysis, AFO, GMB and SMB; Writing & Editing, AFO, GMB and SMB; Supervision, SMB; Funding Acquisition, SMB.

## Supplemental figure legends

**Figure S1. Optimization of FTY720 use in the murine TB model (pertains to Fig.3)**

A. Treatment of B6 mice with FTY720 after Mtb infection led to a reduction in the percentage of T cells (left) and B cells (right) among all live cells in peripheral blood as determined by flow cytometry.

B. Treatment with FTY720 led to significant reductions in the number of total T cells (left) including CD4 (middle) and CD8 (right) T cells in the lungs of Mtb infected B6 mice.

C. The bacterial burden in the lungs of infected B6 mice was increased when treated with 4 mg/kg of FTY720, but not 1 mg/kg of FTY720, as determined four wpi. FTY720 did not have a significant effect on Mtb control in the spleens of treated mice. One-way ANOVA with Dunnett’s multiple comparisons test. Data is from a single experiment. Each point is an individual subject, n=4-5. Box plots indicate median (middle line), 25th, 75th percentile (box) and minimum and maximum (whiskers).

**Figure S2. Gating Scheme for Identifying B & T cells in Peripheral Blood**

Gating strategy for identifying B and T cells in the peripheral blood. In brief, doublets were excluded and then a viability dye was used to exclude dead cells. B cells were identified by expression of CD19 while T cells were identified by expression of CD90 or CD3. Then, CD4 and CD8α or CD8β was used to specifically identify CD4 T and CD8 T Cells.

**Figure S3. Gating Scheme for Identifying T cells in Lung**

Gating strategy for identifying T cell population in the lung. In brief, doublets were excluded and then a viability dye was used to exclude dead cells. B cells were then excluded by expression of CD19, while T cells were identified by expression of CD90. T cells were then subclassified by CD4 or CD8 by their expression of CD4 or CD8α or CD8β, respectively. Naïve T cells were identified by the expression of CD62L and lack of CD44 expression. Antigen-experienced T cells were identified by expression of CD44 and lack of expression of CD62L.

**Figure S4. T cell depletion strategies (pertains to Fig.4)**

A. Evaluation of antibody cross-blocking potential in CC042 mice. The anti-CD4 mAb GK1.5 that is used to deplete CD4 T cells in vivo blocked the binding of labeled GK1.5, but not of alternate anti-CD4 mAbs RM4-4 or RM4-5. The anti-CD8α mAb used in vivo (clone 2.4.3) does not block binding of alternate anti-CD8α clone 53-6.7 or anti-CD8β clone YTS156.7.7. Therefore, the efficiency of CD4 and CD8 T cell depletion in vivo by GK1.5 and 2.4.3 was quantified using anti-CD4 mAbs RM4-4 or RM4-5 and anti-CD8α mAb 53-6.7 or CD8β YTS156.7.7 in all T cell depletion experiments. Similarly, the anti-CD20 mAb MB20-11, used to deplete B cell in vivo, did not inhibit the binding of anti-CD19 mAb clone 6D5 to B cells.
B. Representative flow plot of efficiency of CD4 and CD8 T cell depletion in CC042 mice (left) and quantification of depletion efficiency of total T cells (right top) and CD44+ T cells (right bottom) in the lung, inguinal LN and spleen. Data represents a single experiment. Each point is an individual subject, n=2-3. Line, mean. Error bars, SD. Two-way ANOVA.

**Figure S5. B cell depletion strategies (pertains to Fig.5)**

A. Gating strategy for identifying B cell populations in the lung, lymph node and spleen. In brief, doublets were excluded and then a viability dye was used to exclude dead cells. T cells were excluded by the expression of CD90. Total B cells were identified by the expression of B220. B cells were then further classified as mature or immature. Mature B cells were identified by expression of IgD and lack of CD93 expression, while immature B cells were identified by expression of CD93 and lack of IgD expression.

B. Number of total B cells (top row) and frequency of mature (B220^+^IgD^+^CD93^-–^, bottom row) among total B cells from naïve CC042 mice two weeks in the lung (left), spleen (middle) and iLN (right) after a single administration of anti-CD20 mAb as determined by flow cytometry. Data represents a single experiment. Each point is an individual subject, n=3-4 mice per group. Unpaired t-test. Bars, mean. Line, SD.

**Figure S6. Gating Scheme for Identifying LKS+ Cells in Bone Marrow**

Gating strategy for identifying LKS+ hematopoietic stem cells in the bone marrow. Doublets were excluded and then a viability dye was used to exclude dead cells. LKS+ cells were then identified by the lack of expression of a lineage marker and co-expression of Sca-1 and c-kit. LKS+ cells were then further sub-divided into MPP, ST-HSC and LT-HSC by differential expression of CD48 and CD150.

**Figure S7. Efficacy of antibiotic treatment and combined T cell depletion strategy (pertains to Fig.7)**

A. The efficiency of BCG clearance in the iLN of CC042 mice by the antibiotic treatment regimen is plotted. Representative result from one of two independent experiments. Each dot represents an individual mouse, n=5 mice per group. Paired t-test. The limit of detection was five CFU.

B. Frequency of total CD90^+^ T cells (right), CD4 (middle) or CD8 (left) T cells in peripheral blood as a percentage of total live cells. Data is representative from one of two experiments with similar results using untreated CC042 mice (No Tx), unvaccinated CC042 mice treated with anti-CD4 and anti-CD8α mAbs (Abs/Abx), BCG-vaccinated CC042 mice (BCG) or BCG-vaccinated CC042 mice treated with anti-CD4 and anti-CD8α mAbs (BCG/Abs/Abx). Each point is an individual subject, n=4-5. Bars, mean. Error bars, SD. One-way ANOVA with Šídák’s multiple comparisons test.

C. Frequency of total CD90^+^ T cells (right), CD4 (middle) or CD8 (left) T cells in peripheral blood as a percentage of total live cells. Data is representative from one of two experiments with similar results using unvaccinated B6 mice (B6), unvaccinated CC042 mice (CC042), BCG-vaccinated CC042 mice (BCG) or BCG-vaccinated CC042 mice treated with anti-CD4 and anti-CD8α mAbs (CC042 BCG+T Dep). Each point is an individual subject, n=4-9. Bars, mean. Error bars, SD. One-way ANOVA with Šídák’s multiple comparisons test.

**Figure S8. Histological analysis (pertains to Fig.8)**

A-D. Automated image analysis of histopathological tissue. Three to five lung lobes were analyzed per mouse for a total of 19 (unvaccinated), 27 (BCG), and 34 (BCG+T depletion). **A**. The total area of each scanned lung (left), the total area occupied by lesional tissue (i.e., granulomas, center), and the percentage of the lung occupied by granulomas (right). **B**. Analysis of parameters related to tissue death and destruction including the percentage of lung occupied by cell poor necrosis (left), the percentage of pyknotic debris (middle), and the amount of fibrin (right). **C**. Measurement of parameters relating to the bacterial burden including, from left to right, the number of AFB clusters in debris or in macrophages, and the number of single Mtb in debris or macrophages. **D**. The absolute area of AFB biofilm. Comparisons between different groups were analyzed using a non-parametric one-way Anova (Kruskal-Wallis).

## References

1. World Health O. Global tuberculosis report 2024. Genève, Switzerland: World Health Organization; 2024.

2. Ahmed A, Rakshit S, Adiga V, Dias M, Dwarkanath P, D’Souza G, et al. A century of BCG: Impact on tuberculosis control and beyond. Immunol Rev. 2021;301(1):98–121.

3. Mangtani P, Abubakar I, Ariti C, Beynon R, Pimpin L, Fine PE, et al. Protection by BCG vaccine against tuberculosis: a systematic review of randomized controlled trials. Clin Infect Dis. 2014;58(4):470–80.

4. Fine PE. Variation in protection by BCG: implications of and for heterologous immunity. Lancet. 1995;346(8986):1339-45.

5. Colditz GA, Brewer TF, Berkey CS, Wilson ME, Burdick E, Fineberg HV, et al. Efficacy of BCG vaccine in the prevention of tuberculosis. Meta-analysis of the published literature. Jama. 1994;271(9):698–702.

6. Brosch R, Gordon SV, Garnier T, Eiglmeier K, Frigui W, Valenti P, et al. Genome plasticity of BCG and impact on vaccine efficacy. Proceedings of the National Academy of Sciences. 2007;104(13):5596–601.

7. Coppola M, Jurion F, van den Eeden SJF, Tima HG, Franken K, Geluk A, et al. In-vivo expressed Mycobacterium tuberculosis antigens recognised in three mouse strains after infection and BCG vaccination. NPJ Vaccines. 2021;6(1):81.

8. Moliva JI, Turner J, and Torrelles JB. Immune Responses to Bacillus Calmette-Guérin Vaccination: Why Do They Fail to Protect against Mycobacterium tuberculosis? Front Immunol. 2017;8:407.

9. Churchill GA, Airey DC, Allayee H, Angel JM, Attie AD, Beatty J, et al. The Collaborative Cross, a community resource for the genetic analysis of complex traits. Nat Genet. 2004;36(11):1133–7.

10. Lai R, Gong DN, Williams T, Ogunsola AF, Cavallo K, Lindestam Arlehamn CS, et al. Host genetic background is a barrier to broadly effective vaccine–mediated protection against tuberculosis. The Journal of Clinical Investigation. 2024;133(13).

11. The genome architecture of the Collaborative Cross mouse genetic reference population. Genetics. 2012;190(2):389–401.

12. Smith CM, Proulx MK, Olive AJ, Laddy D, Mishra BB, Moss C, et al. Tuberculosis Susceptibility and Vaccine Protection Are Independently Controlled by Host Genotype. mBio. 2016;7(5).

13. Smith CM, Baker RE, Proulx MK, Mishra BB, Long JE, Park SW, et al. Host-pathogen genetic interactions underlie tuberculosis susceptibility in genetically diverse mice. Elife. 2022;11.

14. Lai R, Williams T, Rakib T, Lee J, and Behar SM. Heterogeneity in lung macrophage control of Mycobacterium tuberculosis is modulated by T cells. Nature Communications. 2024;15(1):5710.

15. Smith CM, Proulx MK, Lai R, Kiritsy MC, Bell TA, Hock P, et al. Functionally Overlapping Variants Control Tuberculosis Susceptibility in Collaborative Cross Mice. mBio. 2019;10(6).

16. Verma NK, and Kelleher D. Not Just an Adhesion Molecule: LFA-1 Contact Tunes the T Lymphocyte Program. J Immunol. 2017;199(4):1213–21.

17. Ley K, Laudanna C, Cybulsky MI, and Nourshargh S. Getting to the site of inflammation: the leukocyte adhesion cascade updated. Nature Reviews Immunology. 2007;7(9):678–89.

18. Masopust D, and Schenkel JM. The integration of T cell migration, differentiation and function. Nature Reviews Immunology. 2013;13(5):309–20.

19. Ghosh S, Chackerian AA, Parker CM, Ballantyne CM, and Behar SM. The LFA-1 adhesion molecule is required for protective immunity during pulmonary Mycobacterium tuberculosis infection. J Immunol. 2006;176(8):4914–22.

20. Sakai S, Kauffman KD, Schenkel JM, McBerry CC, Mayer-Barber KD, Masopust D, et al. Cutting edge: control of Mycobacterium tuberculosis infection by a subset of lung parenchyma-homing CD4 T cells. J Immunol. 2014;192(7):2965–9.

21. Yang Q, Zhang M, Chen Q, Chen W, Wei C, Qiao K, et al. Cutting Edge: Characterization of Human Tissue-Resident Memory T Cells at Different Infection Sites in Patients with Tuberculosis. J Immunol. 2020;204(9):2331–6.

22. Ogongo P, Porterfield JZ, and Leslie A. Lung Tissue Resident Memory T-Cells in the Immune Response to Mycobacterium tuberculosis. Front Immunol. 2019;10:992.

23. Connor LM, Harvie MC, Rich FJ, Quinn KM, Brinkmann V, Le Gros G, et al. A key role for lung-resident memory lymphocytes in protective immune responses after BCG vaccination. Eur J Immunol. 2010;40(9):2482–92.

24. Brinkmann V, Cyster JG, and Hla T. FTY720: Sphingosine 1-Phosphate Receptor-1 in the Control of Lymphocyte Egress and Endothelial Barrier Function. American Journal of Transplantation. 2004;4(7):1019–25.

25. Uchida J, Hamaguchi Y, Oliver JA, Ravetch JV, Poe JC, Haas KM, et al. The innate mononuclear phagocyte network depletes B lymphocytes through Fc receptor-dependent mechanisms during anti-CD20 antibody immunotherapy. J Exp Med. 2004;199(12):1659–69.

26. Garly M-L, Martins CL, Balé C, Baldé MA, Hedegaard KL, Gustafson P, et al. BCG scar and positive tuberculin reaction associated with reduced child mortality in West Africa: A non-specific beneficial effect of BCG? Vaccine. 2003;21(21):2782–90.

27. Netea MG, Domínguez-Andrés J, Barreiro LB, Chavakis T, Divangahi M, Fuchs E, et al. Defining trained immunity and its role in health and disease. Nat Rev Immunol. 2020;20(6):375–88.

28. Butkeviciute E, Jones CE, and Smith SG. Heterologous effects of infant BCG vaccination: potential mechanisms of immunity. Future Microbiol. 2018;13(10):1193–208.

29. Kaufmann E, Sanz J, Dunn JL, Khan N, Mendonça LE, Pacis A, et al. BCG Educates Hematopoietic Stem Cells to Generate Protective Innate Immunity against Tuberculosis. Cell. 2018;172(1):176–90.e19.

30. Dallmann-Sauer M, Fava VM, Malherbe ST, MacDonald CE, Orlova M, Kroon EE, et al. Mycobacterium tuberculosis resisters despite HIV exhibit activated T cells and macrophages in their pulmonary alveoli. J Clin Invest. 2025;135(7).

31. Gideon HP, Hughes TK, Tzouanas CN, Wadsworth MH, 2nd, Tu AA, Gierahn TM, et al. Multimodal profiling of lung granulomas in macaques reveals cellular correlates of tuberculosis control. Immunity. 2022;55(5):827–46.e10.

32. Gela A, Murphy M, Rodo M, Hadley K, Hanekom WA, Boom WH, et al. Effects of BCG vaccination on donor unrestricted T cells in two prospective cohort studies. eBioMedicine. 2022;76.

33. Miranda-Hernandez S, Kumar M, Henderson A, Graham E, Tan X, Taylor J, et al. CD8 (+) T cells mediate vaccination-induced lymphatic containment of latent Mycobacterium tuberculosis infection following immunosuppression, while B cells are dispensable. bioRxiv. 2025.

34. Jung YJ, Ryan L, LaCourse R, and North RJ. Differences in the ability to generate type 1 T helper cells need not determine differences in the ability to resist Mycobacterium tuberculosis infection among mouse strains. J Infect Dis. 2009;199(12):1790–6.

35. Koyuncu D, Tavolara T, Gatti DM, Gower AC, Ginese ML, Kramnik I, et al. B cells in perivascular and peribronchiolar granuloma-associated lymphoid tissue and B-cell signatures identify asymptomatic Mycobacterium tuberculosis lung infection in Diversity Outbred mice. Infect Immun. 2024;92(7):e0026323.

36. Irvine EB, O’Neil A, Darrah PA, Shin S, Choudhary A, Li W, et al. Robust IgM responses following intravenous vaccination with Bacille Calmette-Guérin associate with prevention of Mycobacterium tuberculosis infection in macaques. Nat Immunol. 2021;22(12):1515–23.

37. Divangahi M. Are tolerance and training required to end TB? Nat Rev Immunol. 2018;18(11):661–3.

38. Netea MG, Domínguez-Andrés J, Barreiro LB, Chavakis T, Divangahi M, Fuchs E, et al. Defining trained immunity and its role in health and disease. Nature Reviews Immunology. 2020;20(6):375–88.

39. Netea MG, and Joosten LAB. Trained innate immunity: Concept, nomenclature, and future perspectives. Journal of Allergy and Clinical Immunology. 2024;154(5):1079–84.

40. Mai D, Jahn A, Murray T, Morikubo M, Lim PN, Cervantes MM, et al. Exposure to Mycobacterium remodels alveolar macrophages and the early innate response to Mycobacterium tuberculosis infection. PLOS Pathogens. 2024;20(1):e1011871.

41. Orme IM. Prospects for new vaccines against tuberculosis. Trends in Microbiology. 1995;3(10):401–4.

42. Chackerian AA, and Behar SM. Susceptibility to Mycobacterium tuberculosis: lessons from inbred strains of mice. Tuberculosis (Edinb). 2003;83(5):279–85.

43. Del Pozo-Ramos L, and Kupz A. A review of the efficacy of clinical tuberculosis vaccine candidates in mouse models. Front Immunol. 2025;16:1609136.

44. Henao-Tamayo M, Obregón-Henao A, Creissen E, Shanley C, Orme I, and Ordway DJ. Differential Mycobacterium bovis BCG vaccine-derived efficacy in C3Heb/FeJ and C3H/HeOuJ mice exposed to a clinical strain of Mycobacterium tuberculosis. Clin Vaccine Immunol. 2015;22(1):91–8.

45. Cardona P-J, and Williams A. Experimental animal modelling for TB vaccine development. International Journal of Infectious Diseases. 2017;56:268–73.

46. Zhang J, Teh M, Kim J, Eva MM, Cayrol R, Meade R, et al. A Loss-of-Function Mutation in the Integrin Alpha L (Itgal) Gene Contributes to Susceptibility to Salmonella enterica Serovar Typhimurium Infection in Collaborative Cross Strain CC042. Infect Immun. 2019;88(1).

47. Lee J, Boyce S, Powers J, Baer C, Sassetti CM, and Behar SM. CD11cHi monocyte-derived macrophages are a major cellular compartment infected by Mycobacterium tuberculosis. PLoS Pathog. 2020;16(6):e1008621.

