## Supplemental Figures for "BCG vaccination elicits protection against Mtb infection mediated by two phases of T cell immunity"

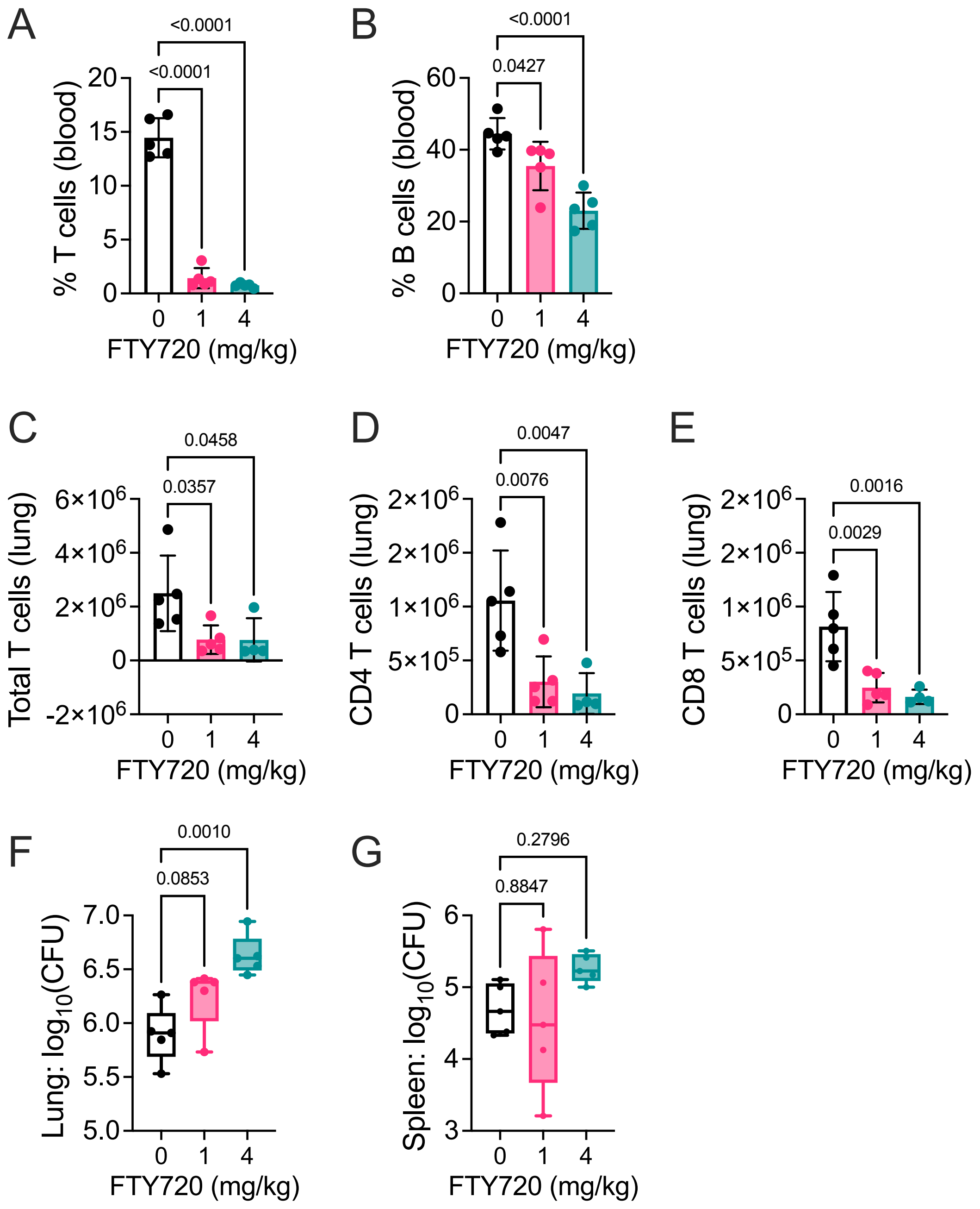


**Figure S1. Optimization of FTY720 use in the murine TB model (pertains to Fig.3)**
A. Treatment of B6 mice with FTY720 after Mtb infection led to a reduction in the percentage of T cells (left) and B cells (right) among all live cells in peripheral blood as determined by flow cytometry.
B. Treatment with FTY720 led to significant reductions in the number of total T cells (left) including CD4 (middle) and CD8 (right) T cells in the lungs of Mtb infected B6 mice.
C. The bacterial burden in the lungs of infected B6 mice was increased when treated with 4 mg/kg of FTY720, but not 1 mg/kg of FTY720, as determined four wpi. FTY720 did not have a significant effect on Mtb control in the spleens of treated mice. One-way ANOVA with Dunnett's multiple comparisons test. Data is from a single experiment. Each point is an individual subject, n=4-5. Box plots indicate median (middle line), 25th, 75th percentile (box) and minimum and maximum (whiskers).

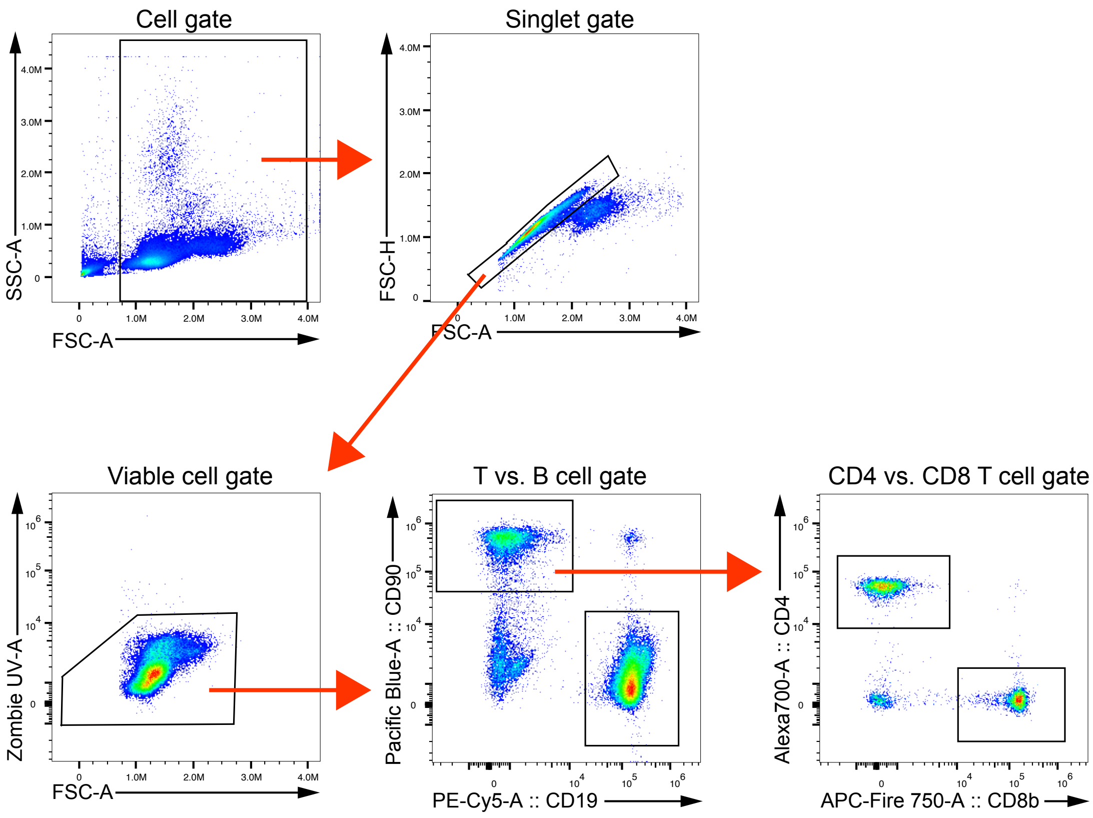


**Figure S2. Gating Scheme for Identifying B & T cells in Peripheral Blood**
Gating strategy for identifying B and T cells in the peripheral blood. In brief, doublets were excluded and then a viability dye was used to exclude dead cells. B cells were identified by expression of CD19 while T cells were identified by expression of CD90 or CD3. Then, CD4 and CD8α or CD8β was used to specifically identify CD4 T and CD8 T Cells.

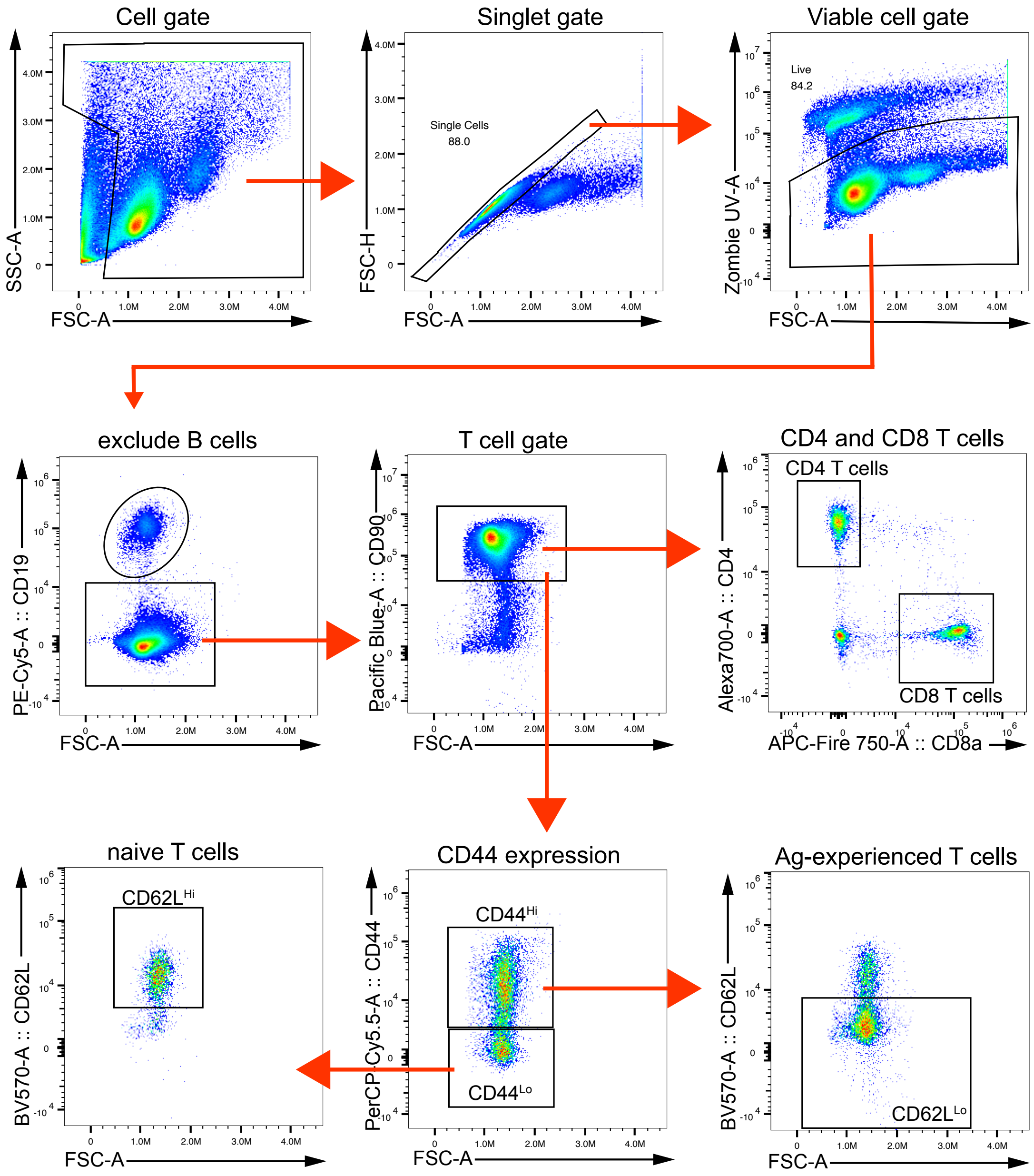


**Figure S3. Gating Scheme for Identifying T cells in Lung**

Gating strategy for identifying T cell population in the lung. In brief, doublets were excluded and then a viability dye was used to exclude dead cells. B cells were then excluded by expression of CD19, while T cells were identified by expression of CD90. T cells were then subclassified by CD4 or CD8 by their expression of CD4 or CD8α or CD8β, respectively. Naïve T cells were identified by the expression of CD62L and lack of CD44 expression. Antigen-experienced T cells were identified by expression of CD44 and lack of expression of CD62L.

| A | Specificity | Primary antibody (in vivo depleting) | Secondary antibody (in vitro staining) | Cross-blocking? |
|---|---|---|---|---|
| | CD4 | GK1.5 | GK1.5 | yes |
| | | | RM4-4 | no |
| | | | RM4-5 | no |
| | CD8a | 2.4.3 | 53-6.7 | no |
| | | | YTS156.7.7 | no |
| | CD20 | MB20-11 | 6D5 | no |

B

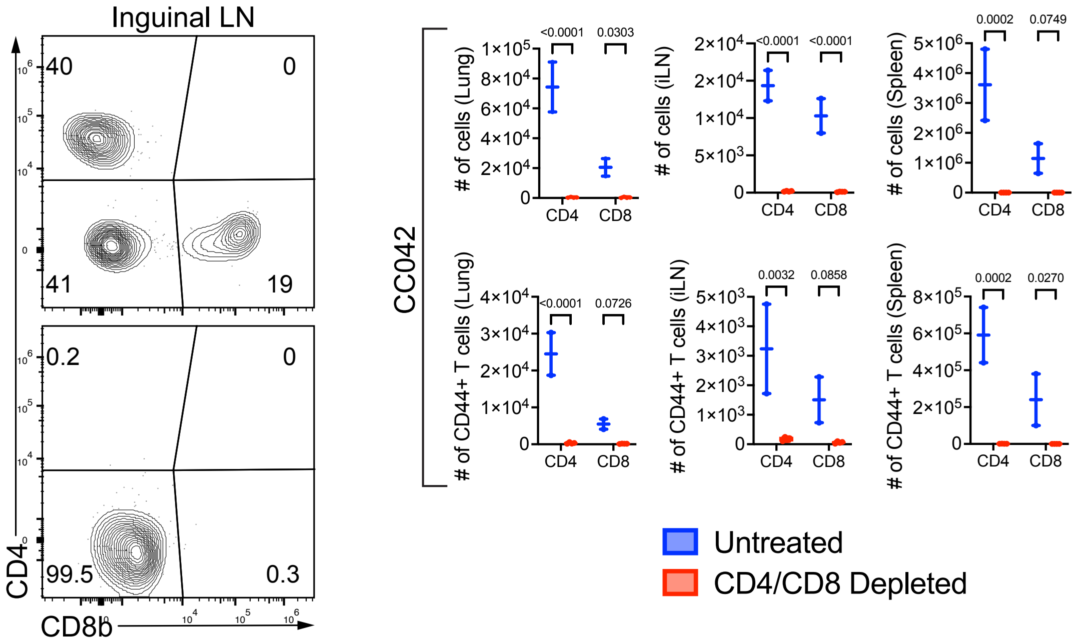


**Figure S4. T cell depletion strategies (pertains to Fig.4)**

A. Evaluation of antibody cross-blocking potential in CC042 mice. The anti-CD4 mAb GK1.5 that is used to deplete CD4 T cells in vivo blocked the binding of labeled GK1.5, but not of alternate anti-CD4 mAbs RM4-4 or RM4-5. The anti-CD8α mAb used in vivo (clone 2.4.3) does not block binding of alternate anti-CD8α clone 53-6.7 or anti-CD8β clone YTS156.7.7. Therefore, the efficiency of CD4 and CD8 T cell depletion in vivo by GK1.5 and 2.4.3 was quantified using anti-CD4 mAbs RM4-4 or RM4-5 and anti-CD8α mAb 53-6.7 or CD8β YTS156.7.7 in all T cell depletion experiments. Similarly, the anti-CD20 mAb MB20-11, used to deplete B cell in vivo, did not inhibit the binding of anti-CD19 mAb clone 6D5 to B cells.

B. Representative flow plot of efficiency of CD4 and CD8 T cell depletion in CC042 mice (left) and quantification of depletion efficiency of total T cells (right top) and CD44+ T cells (right bottom) in the lung, inguinal LN and spleen. Data represents a single experiment. Each point is an individual subject, n=2-3. Line, mean. Error bars, SD. Two-way ANOVA.

A

B

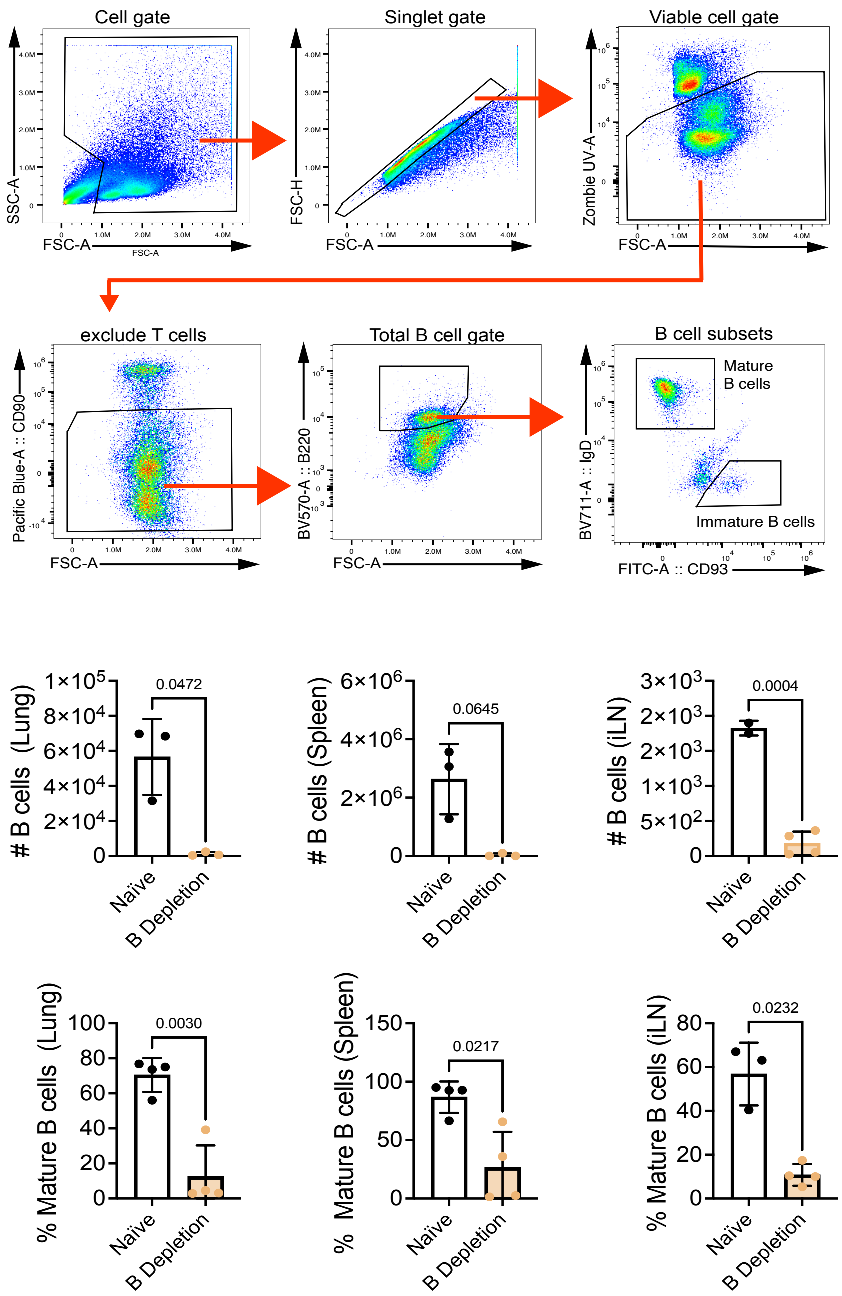


**Figure S5. B cell depletion strategies (pertains to Fig.5).** A. Gating strategy for identifying B cell populations in the lung, lymph node and spleen. In brief, doublets were excluded and then a viability dye was used to exclude dead cells. T cells were excluded by the expression of CD90. Total B cells were identified by the expression of B220. B cells were then further classified as mature or immature. Mature B cells were identified by expression of IgD and lack of CD93 expression, while immature B cells were identified by expression of CD93 and lack of IgD expression. B. Number of total B cells (top row) and frequency of mature ($B220^{+}IgD^{+}CD93^{-}$, bottom row) among total B cells from naïve CC042 mice two weeks in the lung (left), spleen (middle) and iLN (right) after a single administration of anti-CD20 mAb as determined by flow cytometry. Data represents a single experiment. Each point is an individual subject, n=3-4 mice per group. Unpaired t-test. Bars, mean. Line, SD.

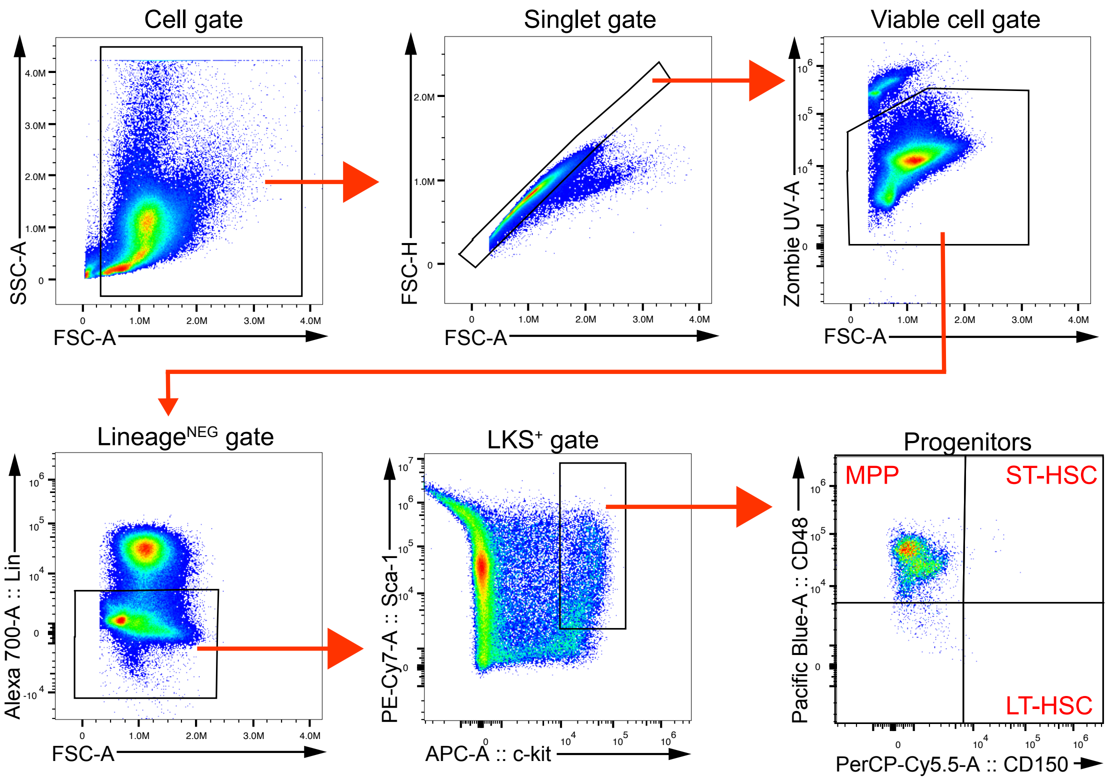


**Figure S6. Gating Scheme for Identifying LKS+ Cells in Bone Marrow**

Gating strategy for identifying LKS+ hematopoietic stem cells in the bone marrow. Doublets were excluded and then a viability dye was used to exclude dead cells. LKS+ cells were then identified by the lack of expression of a lineage marker and co-expression of Sca-1 and c-kit. LKS+ cells were then further sub-divided into MPP, ST-HSC and LT-HSC by differential expression of CD48 and CD150.

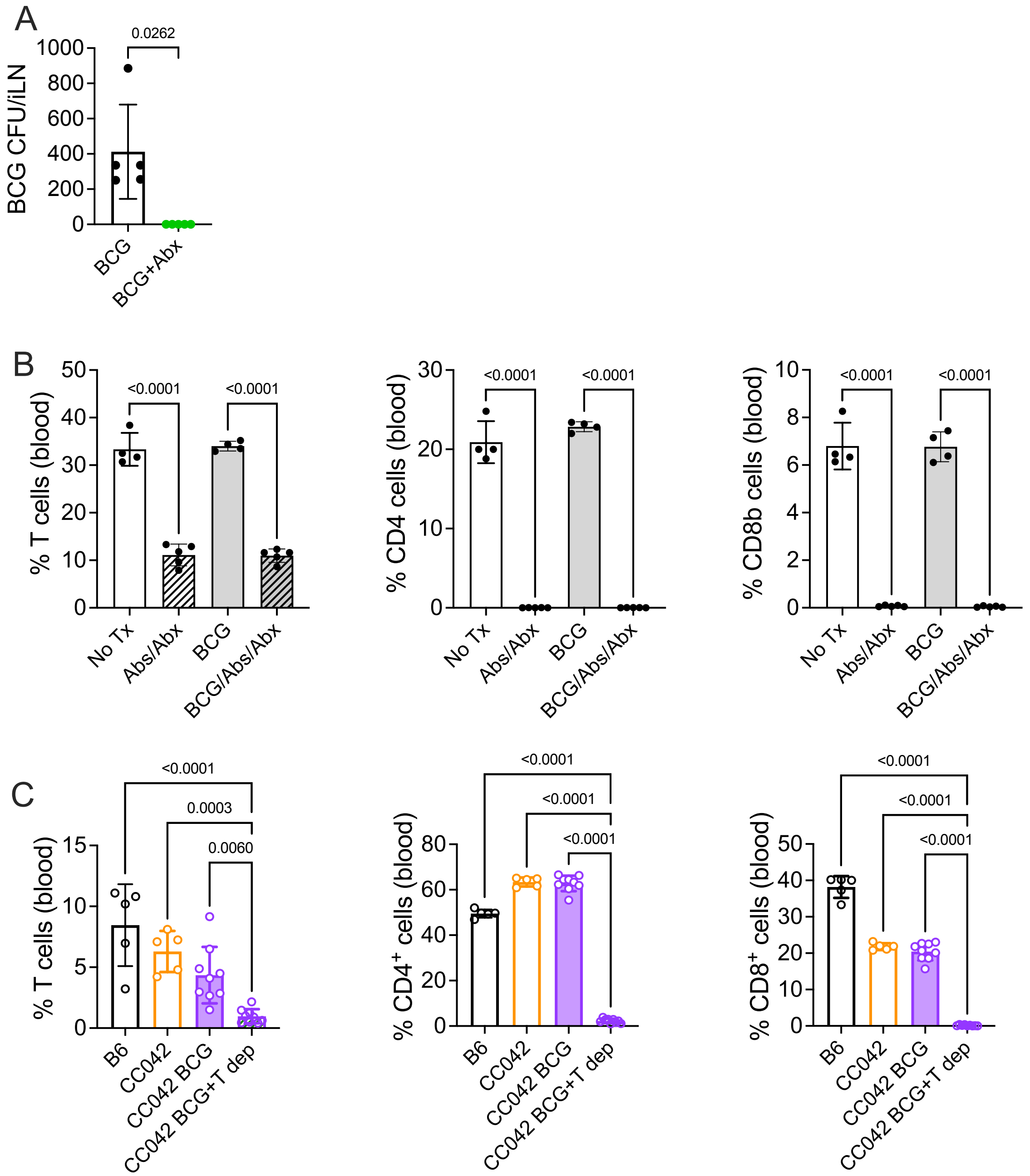


**Figure S7. Efficacy of antibiotic treatment and combined T cell depletion strategy (pertains to Fig.7)**

A. The efficiency of BCG clearance in the iLN of CC042 mice by the antibiotic treatment regimen is plotted. Representative result from one of two independent experiments. Each dot represents an individual mouse, n=5 mice per group. Paired t-test. The limit of detection was five CFU.

B. Frequency of total CD90$^{+}$ T cells (right), CD4 (middle) or CD8 (left) T cells in peripheral blood as a percentage of total live cells. Data is representative from one of two experiments with similar results using untreated CC042 mice (No Tx), unvaccinated CC042 mice treated with anti-CD4 and anti-CD8α mAbs (Abs/Abx), BCG-vaccinated CC042 mice (BCG) or BCG-vaccinated CC042 mice treated with anti-CD4 and anti-CD8α mAbs (BCG/Abs/Abx). Each point is an individual subject, n=4-5. Bars, mean. Error bars, SD. One-way ANOVA with Šídák's multiple comparisons test.

C. Frequency of total CD90+ T cells (right), CD4 (middle) or CD8 (left) T cells in peripheral blood as a percentage of total live cells. Data is representative from one of two experiments with similar results using unvaccinated B6 mice (B6), unvaccinated CC042 mice (CC042), BCG-vaccinated CC042 mice (BCG) or BCG-vaccinated CC042 mice treated with anti-CD4 and anti-CD8α mAbs (CC042 BCG+T Dep). Each point is an individual subject, n=4-9. Bars, mean. Error bars, SD. One-way ANOVA with Šídák's multiple comparisons test.

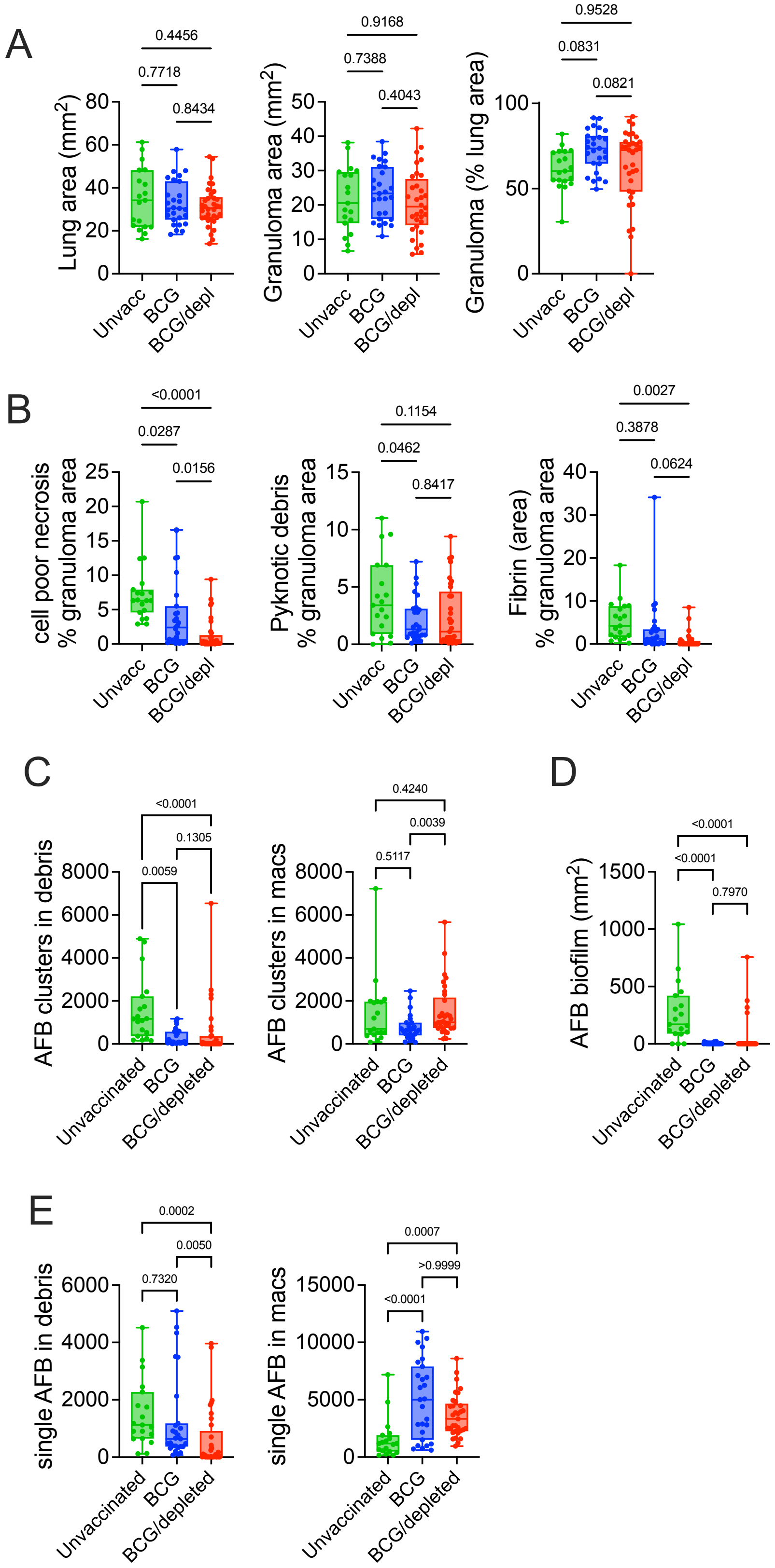

A
Lung area (mm2)
Granuloma area (mm2)
Granuloma (% lung area)
Unvacc
BCG
BCG/depl
0.4456
0.7718
0.8434
0.9168
0.7388
0.4043
0.9528
0.0831
0.0821
B
cell poor necrosis % granuloma area
Pyknotic debris % granuloma area
Fibrin (area) % granuloma area
<0.0001
0.0287
0.0156
0.1154
0.0462
0.8417
0.0027
0.3878
0.0624
C
AFB clusters in debris
AFB clusters in macs
Unvaccinated
BCG
BCG/depleted
<0.0001
0.1305
0.0059
0.4240
0.0039
0.5117
D
AFB biofilm (mm2)
<0.0001
<0.0001
0.7970
E
single AFB in debris
single AFB in macs
0.0002
0.0050
0.7320
0.0007
>0.9999
<0.0001

**Figure S8. Histological analysis (pertains to Fig.8)**

A-D. Automated image analysis of histopathological tissue**.** Three to five lung lobes were analyzed per mouse for a total of 19 (unvaccinated), 27 (BCG), and 34 (BCG+T depletion). **A**. The total area of each scanned lung (left), the total area occupied by lesional tissue (i.e., granulomas, center), and the percentage of the lung occupied by granulomas (right). **B**. Analysis of parameters related to tissue death and destruction including the percentage of lung occupied by cell poor necrosis (left), the percentage of pyknotic debris (middle), and the amount of fibrin (right). **C**. Measurement of parameters relating to the bacterial burden including, from left to right, the number of AFB clusters in debris or in macrophages, and the number of single Mtb in debris or macrophages. **D**. The absolute area of AFB biofilm. Comparisons between different groups were analyzed using a non-parametric one-way Anova (Kruskal-Wallis).
